# Origins of the Fittest: Clonal Interference in Heterogeneous Networks

**DOI:** 10.1101/2025.10.20.683208

**Authors:** Adrian Zachariae, Pascal Klamser, Dirk Brockmann

## Abstract

In asexual populations, clonal interference, the competition between strains carrying different beneficial mutations, plays a crucial role in shaping evolutionary outcomes. This study investigates how this phenomenon unfolds in complex heterogeneous networks, demonstrating that network structure significantly affects the origin and spread of high-fitness strains.

Using computational modeling and a novel analytical approach, we demonstrate that the interplay between mutation rates and network topology creates distinct evolutionary regimes. We observe that the mutations driving adaptation originate from network locations that facilitate mutation spread within a time-frame given by the mutation rate. If the window widens or narrows, the contribution of different network regions grows or diminishes. Using analytical approaches from epidemic modeling and network geometry, our analysis efficiently captures this evolutionary dynamic in arbitrary network configurations.

This work provides new insights into how spatial heterogeneity shapes evolutionary trajectories and indicates that attempts to alter the speed of adaptation through network modifications will succeed or fail depending critically on the prevailing evolutionary regime—insights relevant to epidemiology, conservation biology, and beyond.

## I. INTRODUCTION

The process of adaptation, or the ability of a population to evolve and adapt to its environment, is a key area of study in evolutionary biology. In the *strong-selection weak-mutation regime*, the traditional or naïve notion of adaptation is that rare mutations accumulate sequentially [1–4]. Here, the speed of adaptation depends linearly on both the mutation rate and the fixation probability. However, for asexual populations, if the population size and mutation rates are large enough, the phenomenon known as clonal interference (CI) disrupts these linear relationships. Clonal interference was first proposed in [3] and has since been observed in various experimental studies [5–7]. CI occurs when mutational events are so frequent that new mutations often occur before previous mutations have fixed, in the so-called *multiple-mutation regime* [8, 9](sometimes called the *concurrent mutation regime* [10] or *clonal-interference regime* [11]). Without the ability of sexual populations to recombine, adaptive mutations must compete on separate evolutionary strands, resulting in a slowing of adaptation. This phenomenon was originally described for well-mixed, unstructured populations [3].

However, it is well recognized that the topological structure of a population can have a large influence on the outcome of evolutionary processes. In the field of evolutionary graph theory, the influence of complex population structures on fixation probability and fixation time has been studied extensively [12–17] . While fixation probability is the main determinant of adaptation speed in the *strong-selection, weak-mutation regime*, structures that increase fixation probability often increase fixation time and may suppress adaptation at higher mutation rates [18–20]

Previous work has studied clonal interference in heterogeneous metapopulation structures [8, 9, 21]. The models used, such as the Wright-Fisher model [22–25] make a sharp distinction between well-mixed population dynamics within subpopulations and migration rates between subpopulations. They have generally found that slow migration rates increase clonal interference. Some works have developed theoretical frameworks to accurately quantify the rate of adaptation in selected network configurations [21]. The meta-populations in these studies have generally featured simple network configurations between subpopulations, such as pairs, rings, or fully connected networks [21] or have featured the random linkage of an Erdős-Rényi model [8, 9]. The focus of these studies has been on population-level outcomes, such as the rate of adaptation, and primarily relied on simulations [8, 9, 26].

Clonal interference has also been studied in the context of structured populations. In particular, lattice-like homogeneous population structures have been studied extensively [11, 27–29]. A key takeaway from such studies is that the absence of long-range connections or long-range dispersal in lattice-like structures accentuates CI, further slowing adaptation.

Some of these studies have noted the similarities between the spread of beneficial mutations and waves or other physical spreading phenomena, and used this analogy to motivate their analysis [11]. Real-world population structures of many asexual organisms, such as pathogens spreading through host social networks, often have highly heterogeneous structures, with large variations in clustering and degree and hierarchical organization. In the fields of epidemics and network geometry, much effort has been devoted to understanding how dynamical processes spread through such complex networks. Some remarkably simple approaches using distances defined along shortest or random paths [30, 31] or differential equations of only pairwise coupled nodes [32–34] have been very successful in approximating various spreading dynamics such as disease dynamics, opinion dynamics, or signal propagation [35–38].

In this paper, we pursue two complementary goals. First, we adapt methods originally developed to analyze spreading phenomena in complex networks to the study of clonal interference in heterogeneous environments. Second, we shift the focus from global, population-level outcomes to the relative contributions of individual locations within the network to the overall speed of adaptation.

Our investigations show that heterogeneous network structures can drastically change the spreading dynamics of new strains and with them the speed of adaptation, sometimes exceeding the expected speed of adaptation in equivalent well-mixed populations.

They also introduce a new aspect to the CI dynamics not seen in homogeneous habitats: How the origin location of new lineages relates to their relative fitness and success. In heterogeneous structured populations, a mutation’s fate is not determined solely by its fitness advantage but also by the position of its origin node within the network. Mutations that arise in regions that facilitate spread across the network are more likely to succeed, linking evolutionary success to network centrality. This coupling between lineage origin and evolutionary outcome represents a fundamental difference from the dynamics in homogeneous populations.

Due to the inherent non-linearity of CI, the interplay between the spreading rates within the population and the mutation rate is complex and can generate several distinct regimes. Interestingly, the most likely origin of successful lineages can change completely between these regimes. These shifts follow a simple rule: The fittest lineages originate from nodes that facilitate the spread of the mutation within the time-frame given by the mutation rate. Each mutation rate thus corresponds to a specific scale in the network. For slow mutation rates, nodes that are central to the network on a global scale are the most likely origins. For faster mutation rates, the scale becomes progressively smaller until only very local node attributes like the node degree dominate.

As the relative contribution of subnetworks to the speed of adaptation changes, modifications to the network aimed at curbing the speed of adaptation will vary in their relative effectiveness based on the mutation rate.

Our results are supported by an analytical approach that focuses on the fittest strains. This approach, which separates propagation dynamics and transitions between strain origins, remains feasible for larger systems. We also show how this approach can be simplified drastically by using distances (i.e. single values) to represent the spreading rates between network locations, while still predicting the qualitative regimes of each network. Understanding clonal interference in heterogeneous environments has important practical implications. Network structure shapes the speed of adaptation of clonal organisms, which in turn affects our understanding of evolutionary timescales. But it also affects the individual contributions to the adaptation process. Insights into how mutations spread and compete can therefore inform strategies to manage or slow adaptation, for example in the context of pathogen evolution.

## II. MODEL DEFINITION

The habitat is represented as a weighted network consisting of *N* nodes and *M* links. Each node represents a sub-population of identical size. The dynamics of that sub-population are assumed to be fast compared to the inter-site dynamics.

One natural example of such a scenario would be an obligatory symbiotic organism inhabiting a single host species. Here, the network represents the social network of *N* host organisms. Another example would be a habitat fractured by human infrastructure or agriculture.

Each node records the dominant strain in the corresponding organism sub-population and each strain *v* is solely characterized by its fitness *r*_*v*_.

A site’s resident strain, *v*, can change in two ways:

1. Through mutation within the resident clone population at node *i*.
2. Through replacement by a fitter strain *u* originating from a neighboring node *j*.

We assume that the subpopulations are large enough that the fixation of deleterious mutations in a subpopulation can be neglected. We do not consider the effects of epistasis and assume an infinite sites model, where mutations accumulate infinitely and at a constant rate. Furthermore, we assume that neutral or disadvantageous mutations do not become dominant within their sub-population and that each mutation event increases fitness.

Based on these assumptions, mutation events can be modeled by a Poisson process describing the combined probability that a new mutation occurs and survives random drift. This process is characterized by a uniform rate *µ*, which is a function of the sub-population of each node, the per-genome mutation rate and the probability that the mutation survives random drift. Such an event in site *i*, whose resident strain is *u*, results in the creation of a new strain *w* with the following rule:

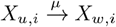

assuming a constant fitness increment *s* = *r*_*w*_ − *r*_*u*_, where *r*_*u*_ and *r*_*w*_ are the fitness values of strain *u* and *w*, respectively (see appendix F 1 for results with an exponentially distributed fitness increase).

Other than by mutation, strains can also be replaced by invasion of a fitter strain from a neighboring node.

The node *j* is a neighbor of node *i* if the element *A*_*ij*_ of the weighted adjacency matrix *A* is nonzero. We restrict our analysis to symmetric networks, *A*_*ij*_ = *A*_*ji*_.

Invasions between neighbors occur at a constant rate given by the link weights *A*_*ij*_ and succeed based on the probability of the invading strain to fixate (≈ 2*s*, assuming a well-mixed population [39]). We consider the fixation time of an invading strain to be short compared to the rate of invasion events and thus treat each invasion event as instantaneous.

The process representing invasion events from *j* → *i* that replace the resident clone of strain *v* with a clone of the invading strain *u* is analogous to the SI epidemic model and can be represented by the following local rule:

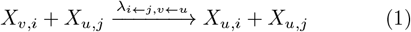

The rate *λ*_*i*←*j,v*←*u*_ depends linearly on both the selective fitness advantage *r*_*u*_ − *r*_*v*_ and the link weight *A*_*ij*_:

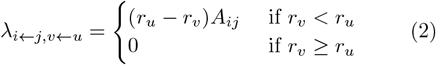

This formulation of invasion events mirrors SI infection dynamics and in a system with just two strains and in the absence of mutation events the dynamics are identical.

The model operates in continuous time to accommodate systems with highly heterogeneous rates. Each simulation uses Gillespie’s Stochastic Simulation Algorithm (SSA), which permits a mathematically exact implementation of the rules above [40]. The SSA and its application to our model is detailed in Appendix B and illustrated in Fig. A1.

## III. RESULTS

### A. Scaling of adaptive speed with network size in different networks

Small-world networks are a class of networks characterized by short distances between nodes: The shortest path between any two randomly chosen nodes in the network only includes a small number of links, *L*, that increases logarithmically with the size *N* of the network:

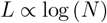

This property is very common in real-world networks, including social networks, and can be easily replicated by a variety of simple model networks.

Among the various network models that exhibit the small-world property, one of the simplest and most fundamental is the Erdős–Rényi (ER) network model [41–43]. In networks built with the ER model, each possible link between two nodes is equally likely and each node in an ER network has a binomially distributed number of neighbors, with only limited variability.

Many real networks have a much higher variability in node degree. This led to the development of other network models that better represent heterogeneous link distributions. One of the simplest and best known networks of this class is the Barabási–Albert (BA) network [44].

The link distribution in the BA network is approximately scale-free, a property that also characterizes many natural networks.

We compare the adaptation rate in our system on ER and BA networks with two well-studied population configurations: The well-mixed population and a planar grid. To ensure that the population structures are comparable, we keep the average weighted degree of each node constant across networks and normalize it to 1 by assigning each link weight (*A*_*ij*_ = *a*) the value *a* = 1*/* ⟨*k*⟩, the inverse of the unweighted degree. This ensures that the overall rate of invasion events is identical across networks of the same size. Between the ER, BA and planar grid networks we also keep the number of links the same at ⟨*k*⟩ = 8. These networks have the exact same number of nodes and links (of identical link strength), and just differ in the wiring pattern. Additional details about the construction of each network can be found in Appendix A.

Fig. 1 shows the adaptive rate 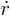 for all four population configurations: As expected the adaptive rate saturates quickly with *N* in the grid configuration to a constant maximum [11] and the well-mixed configuration shows a transition from linear to logarithmic growth with *N* [5, 10].

**FIG. 1.**
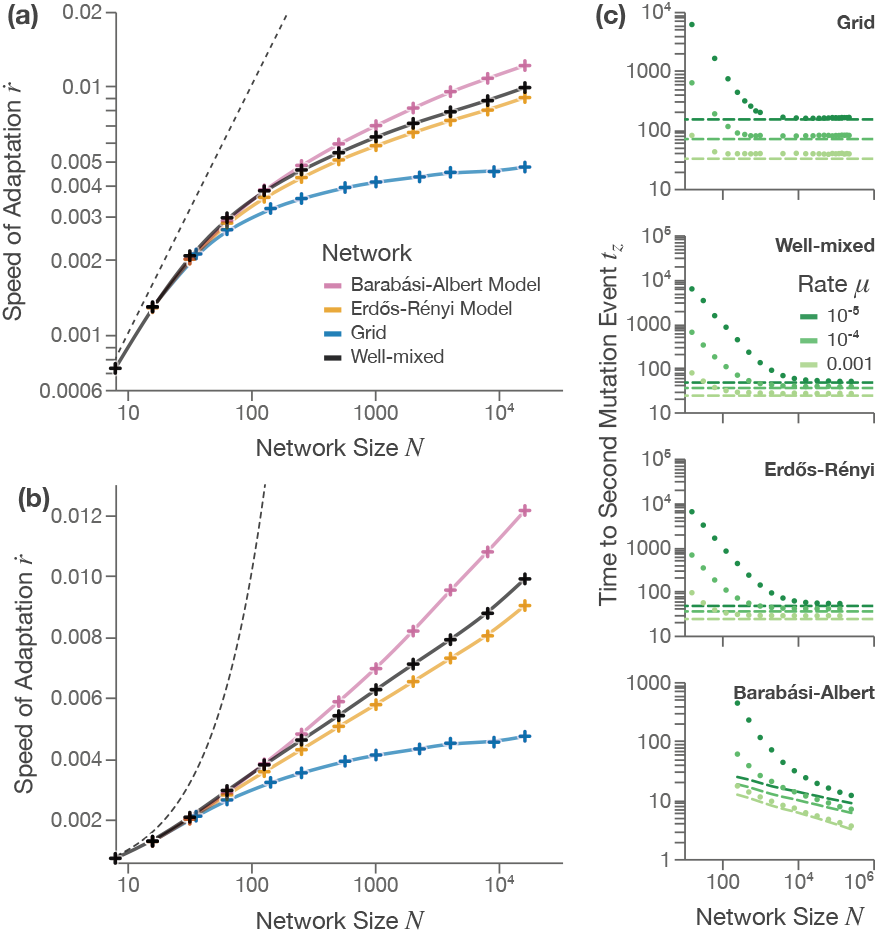
CI in classical small-world network models versus established well-mixed and planar habitat configurations. (a, b): The scaling behavior of the speed of adaptation 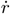 with the habitat size *N* . For comparison the total rate of mutations *µN* is shown (black dotted line). Simulations run with a simulation time of *T/*(*µN*) = 10^5^ and *s* = 0.1. (c): The time *t*_*z*_ until successive accumulation of mutations for different muta-tion rates *µ* and habitat sizes *N* . *t*_*z*_ denotes the time it takes for a strain *y* spreading through an otherwise homogeneous population of lower fitness (*x*, with *r*_*y*_ = *r*_*x*_ + *s*) to produce a strain *z* with another mutation and *r*_*z*_ = *r*_*y*_ + *s* = *r*_*x*_ + 2*s*. Dots represent simulations of *t*_*z*_ (see Appendix B 5 a), lines the approximate solutions from eq. 8, 9, 13 (for⟨ *t*_*z*_ ⟩^grid^, with *c* ≈ 0.063 estimated from simulations). For comparability, the average weighted degree of the habitat networks is normalized to ⟨*k*^*w*^⟩ = 1, and the Grid, ER, BA networks have the same number of links, *M* = 4*N* . Data Points represent averages over 1,000 realizations.

The adaptive rates in the ER network configurations are very close, but slightly slower than, the well-mixed network and also show an approximately logarithmic scaling with *N* for large *N* .

The BA network however, shows the smallest slowdown and stays above logarithmic scaling with *N* (Fig. 1 a.

Our findings agree well with the notion of BA networks as “ultra-small-world networks” that are known to facilitate spreading beyond equivalent well-mixed configurations [45]. Just as the transition from a planar to a small-world facilitates the spread of mutations and reduces clonal interference, the additional speed-up from Small-world configurations to ultra-small-world further decreases CI.

### B. The spreading dynamics of two consecutive mutations

In our simple model system, any slowdown in the rate of adaptation 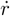 below the rate of all mutations *µN* must result from mutations occurring in strains of lower fitness than the current maximum.

Thus, we find it useful to distinguish between mutations that create a strain with a fitness *s* higher than the fitness of all strains present in the population, and those that do not. We call mutation events that produce such a strain *adaptive*. We denote the average waiting time between adaptive mutation events by ⟨*τ* ⟩.

In finite and fully connected populations with a constant fitness increment *s*, the inverse of ⟨*τ* ⟩ has a simple linear relationship with the rate of adaptation 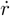:

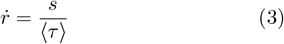

For moderate mutation rates, the spread of the current fittest strain typically outpaces the generation of new mutants. In this regime, the growth of *y* is dominated by invasion events, and the contribution of additional mutation events can be neglected. This approximation is standard in analyses of clonal interference [10, 35] and becomes increasingly accurate in the full system, where the clone that produced *y* itself occupies only a fraction of all nodes (Appendix D 1).

Consequently, ⟨*τ*⟩ depends mainly on the spreading dynamics of the strains that are produced by an adaptive mutation and that determine how quickly the strain can produce a new strain with another successive mutation. The spreading dynamics of the fittest clone depends on both the habitat network and on the fitness difference of that clone to the other clones that it encounters. We first consider a simplified case, that focuses entirely on the network structure.

Starting from a completely homogeneous population with fitness *r*_*x*_, we introduce a single mutation that creates a clone with the fitness *r*_*y*_ = *r*_*x*_ + *s*, creating a strain of fitter organisms spreading through the habitat. We call the strain of the growing population *y* and its abundance *Y* (*t*). At any time *t*, the population *Y* (*t*) generates new established mutants with a rate *µY* (*t*).

We are primarily interested in the event in which the first established mutant arises from *y*, an event we denote with *y* → *z*. We denote the waiting time for *y* → *z* with *t*_*z*_ and its average with ⟨*t*_*z*_⟩ .

Note that in our simple CI model, the spreading dynamics of *y* up until the *y* → *z* event are identical to the spreading dynamics of an SI epidemic compartment model.

To estimate ⟨ *t*_*z*_⟩, we consider the probability density *f* (*y* → *z, t*) of the mutation event *y* → *z*. Starting with the discretized probabilities, the probability *P* (*y* → *z, i*Δ*t*) that the first *y* → *z* event occurs in the *i*-th interval Δ*t*:

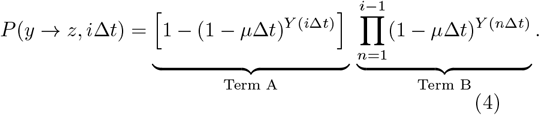

Here, Term A is the probability of at least one *y* → *z* mutation event in the *i*-th interval and Term B the probability of no *y* → *z* mutation event before the *i*-th interval. For small Δ*t*, (1 − *µ*Δ*t*)^*Y* (*n*Δ*t*)^ ≈ exp(−*µY* (*n*Δ*t*)Δ*t*), so if Δ*t* → 0, the CDF and PDF are:

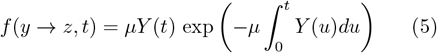

In principle *Y* (*t*) is a random variable, but when its coefficient of variation is small, we may replace it by its deterministic mean. This is usually appropriate for most structures if growth of *y* outpaces the mutation rate (see Appendix D 2 for ta comparison using the three network models from section III A). As the distributions are strictly positive, the mean of the distributions can be computed via the survival function *S*(*y* → *z, t*) = 1 − *F* (*y* → *z, t*):

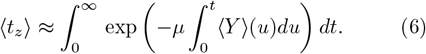

Note that the dynamics of *y* depend linearly on the parameter *s*. That means that a scaled version of the spreading dynamics can be used to separate the structural component (described by ⟨*Y* ^*^⟩ (*t*) with *s* = 1) and the biological parameters (*µ* and *s*), by substitut-ing 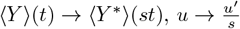 and 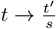:

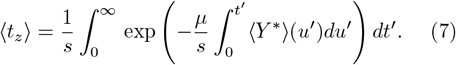

To progress further, we need ⟨*Y* ^*^⟩ (*t*), which depends on the habitat network structure. First, we will show how this equation reduces to well-known formulas in the case of the two homogeneous habitat structures, the planar grid network and the well-mixed habitat.

For the planar grid (⟨*t*_*z*_⟩ ^grid^), we approximate ⟨*Y* ^*^ ⟩ (*t*) with a circular wave pattern of speed *c* ∝ *a*^2^ (*a* is the homogeneous link weight): ⟨*Y* ^*^⟩ (*t*) = *cπt*^2^. For the well-mixed populations (⟨*t*_*z*_⟩ ^WM^), we use the exponential growth function ⟨*Y* ^*^ ⟩ ≈ *e*^*at*^. Both equations have analytical solutions:

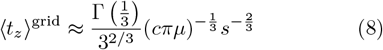

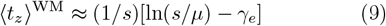

where *γ*_*e*_ = 0.57721 … is the Euler–Mascheroni constant. Eq. 8 is very similar to the scaling law that Martens and Hallatschek [11] found for the speed of adaptation in planar habitats and shows the same scaling behavior with *µ* (of 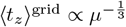), while Eq. 9 is very close to the approximate formula of ⟨*t*_*z*_⟩ = (1*/s*)[ln(*s/µ*) − 1] in [46]. These formulas for ⟨*Y* ^*^ ⟩ (*t*) only consider the unbounded growth of *y* before the wave is able to cover the whole habitat and only become accurate for large enough *N* (see Fig. 1c).

How can we apply this approach to the ER and BA networks? While the exact solution of epidemic spreading dynamics on networks are themselves a high-dimensional problem, we can draw from the vast prior work exploring epidemic dynamics of networks to find suitable approxi-mations for ⟨*Y* ^*^⟩(*t*).

Here, we use an approximate solution of ⟨*Y* ^*^⟩(*t*) that is based on the heterogeneous mean-field approximation, HMF, also called the degree-based mean-field approximation [47].

This approximation has been specifically developed to capture the effect of link heterogeneity on spreading dynamics. It treats all nodes with the same degree *k* as statistically equivalent, but allows the “infected” fraction of nodes to differ between nodes with different degrees. Since nodes with many links both spread the contagion faster and are easier to reach, the high-degree compartments see accelerated early growth after which the contagion cascades to lower-degree nodes [48, 49]. For small *t*, where *Y* (*t*) ≪ *N*, and for uncorrelated connections between nodes of differing degrees, the approximate solution for the spreading dynamics is well known:

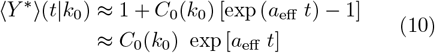

with 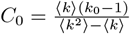 and 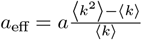. The variable *k*_0_ denotes the degree of the node that the strain *y* orig-inated from, ⟨*k*⟩, and ⟨*k*^2^ ⟩the first and second moment of the degree distribution. With increased heterogeneity, the ratio ⟨*k*^2^ ⟩*/* ⟨*k*⟩ increases and the exponential term of the equation increases more steeply.

Eq. 10 is structurally similar to the well-mixed case (of ⟨*Y* ^*^⟩ (*t*) ≈ *e*^*at*^, detailed in eq. 9, and by analogy we derive:

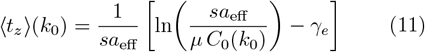

Because of the importance of the heterogeneity ratio ⟨*k*^2^⟩ */*⟨ *k*⟩ to epidemic spreading processes, it is well established for fundamental network models such as ER and BA.

In the ER network, ⟨*k*^2^⟩ = ⟨*k* ^2^⟩ + ⟨*k*⟩, independent of *N*, and while in general ⟨*t*_*z*_⟩ depends on *k*_0_, the total variance is small, in the order of ≈ 1*/*(*sa*_eff_ ⟨*k* ⟩) (Appendix D 3).

Thus, we can substitute ⟨*t*_*z*_⟩(*k*_0_) with:

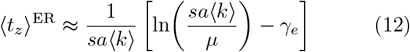

With our normalization of *A* of *A*_*ij*_ = *a* = 1*/*⟨ *k* ⟩, we recover the exponential of the well-mixed case: ⟨ *t*_*z*_⟩ ^ER^ ≈ (1*/s*)[ln(*s/µ*) − *γ*_*e*_], see Fig. 1c.

However, for the BA habitat network, the heterogeneity ratio ⟨*k*^2^ ⟩*/* ⟨*k*⟩ diverges for *N* → ∞, with ⟨*k*⟩ = 2*m* and ⟨*k*^2^⟩ ≈ 2*m*^2^ ln(*N/m*), where *m* is the number of links added with one node per construction step for the BA network (see Section A).

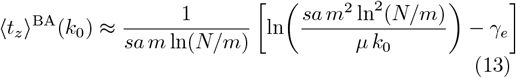

This means that for increasing *N* the waiting time ⟨*t*_*z*_⟩ ^BA^ decreases for all degree classes of nodes, particularly for the nodes with the highest degrees in the network (see Fig. 1c). Note that because of the high variance in *k*, we cannot easily ignore the origin degree *k*_0_.

#### 1. Comment on fitness variance

There is an important distinction between the waiting time ⟨*t*_*z*_⟩ and the waiting time between adaptive mutations in the full model, ⟨*τ* ⟩.

In the full model a new mutation will not encounter a homogeneous population, but instead a mixture of other strains with a distribution of fitness values. The exact distribution of fitness values encountered by a spreading mutation will in general depend on the exact location of the previous mutation events and it’s very hard to quantify precisely. However, we find that a mean field approximation that uses a homogeneous average fitness difference ⟨*s*_Δ_⟩, gives satisfactory results if ⟨*Y* ⟩(*t*) is not very small in the time-frame of ⟨*τ* ⟩ (as seen in appendix G using sampled ⟨*s*_Δ_⟩-values).

By substituting *s* with *s*_eff_ in Eq. 6, we arrive at the equation:

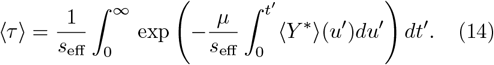

The new dynamics of *y* have strong parallels to epidemic spreading dynamics in populations with heterogeneous susceptibility, a problem as described in [50, 51]. In sparse networks, if high susceptibility (analogous to large fitness differences) is anti-correlated with centrality (as to be expected in our case), this will slow down spread compared to the homogeneous case [51]. As such the *s s*_eff_ *s*_Δ_, where *s*_Δ_ is the average fitness difference of the fittest strains to all other nodes. These boundaries can be though of as the two extreme cases: The first is a habitat with a grid structure and of sufficient size (relative to *µ/*(*s a*)). Here, any mutation will spread in a predictable circle-pattern with little stochasticity and mutations (in the fittest lineages) will very likely only ever encounter their direct predecessor clone until the next mutation event. This means *s*_eff_ *s* and *τ t*_*z*_, and *s*_Δ_ becomes inconsequential, as consistent with [11]. The other is the well-mixed system, in which no structure can induce a bias in the strains encountered and *s*_eff_ = *s*_Δ_ .

In analogy to the approach for well-mixed populations in [10, 46], the quantities ⟨*s*_Δ_⟩ and 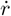 can be connected via the arrival time 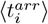 of a new adaptive mutation to a node *i*: the average fitness increase in *any* node via both mutation and (predominantly) invasion events needs to match 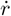. For most scenarios this will be dominated by the invasion events. We can then estimate *s*_eff_ by matching the two rates (see appendix G).

For the small networks in the following sections, ⟨*s*_Δ_⟩ will not be much greater than *s* up until *µ* ≈ *sa* and we will use *s*_eff_ ≈ *s*.

### C A model of many consecutive mutations in networks

There is another complication in CI in heterogeneously structured populations that is absent in homogeneous populations. In homogeneous environments, there is no distinction between the *x* → *z* spreading dynamics originating from different locations. For heterogeneous networks, the evolution of *Y* (*t*) depends on the origin node *j* of the strain *y*. By extension, the average waiting time ⟨*τ*_*j*_ ⟩ also depends on *j*, as shown for BA networks in Eq. 13, where ⟨*t*_*z*_⟩ ^*BA*^(*k*_0_) depends on the degree of the origin node *k*_0_ (illustrated in Fig. 2a). Additionally, the origin nodes of the fittest lineages might also be of intrinsic interest to researchers. Here, we present a framework to quantify the distribution *π*, which denotes the relative frequency of adaptive mutation events in each node, i.e. mutation events that produce mutants with a higher fitness than all other individuals in the population.

**FIG. 2.**
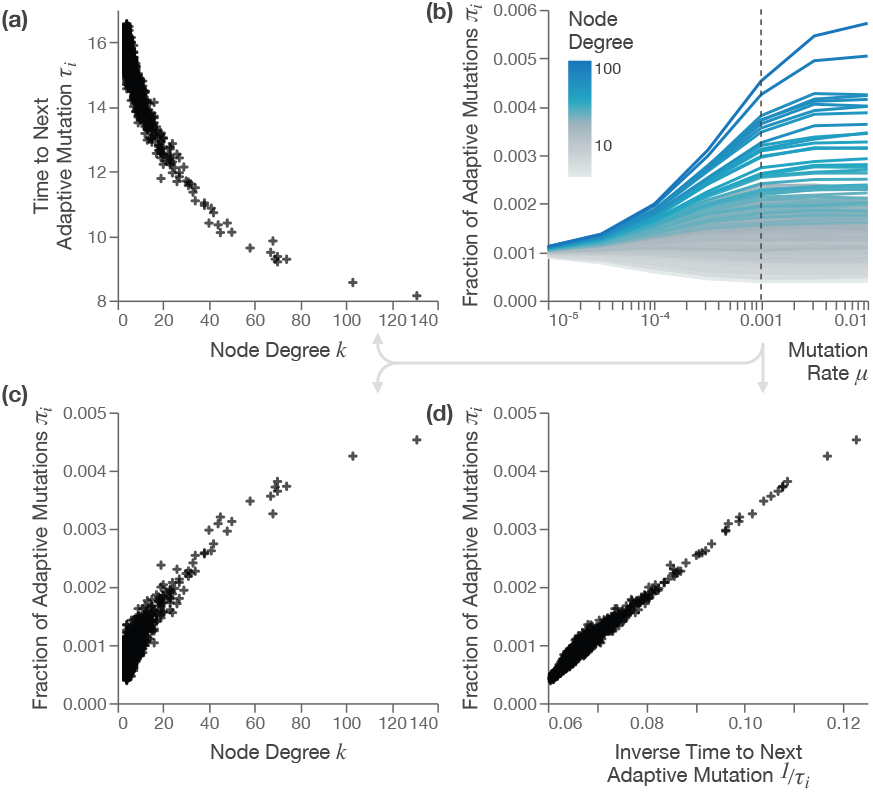
Dependence of average waiting time between mutations, ⟨*τ*_*i*_⟩, and of the fraction of adaptive mutations originating in a node, *π*_*i*_ in BA networks. (a): The node degree *k* dependence of ⟨*τ*_*i*_ ⟩ for *µ* = 0.001. (b): As the mutation rate *µ* increases, the distribution of *π*_*i*_ widens. (c): The node degree *k* dependence of ⟨*π*_*i*_⟩ for *µ* = 0.001. (d): As predicted by Eq. 21, the relationship between ⟨*τ*_*i*_ ⟩ and *π*_*i*_ is near-linear (shown for *µ* = 0.001). All figures show the same realization of a BA network with *N* = 1024. Values are averages over 10,000 simulation trajectories with a simulation time of *T* = 16*/µ* and *s* = 0.1.

Naively, one might assume that the density of origin nodes of the fittest lineages is just homogeneous, so that we could compute ⟨*τ*⟩ (and by extension 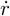) by simply averaging over all possible origins *j*. However, if we look at the simulation data on the BA-network structure, the origins are clearly not equally distributed within the CI-regime (see Fig. 2b-c). We observe a strong dependence of the origin of a new fittest strain on the node degree in BA-like networks. This makes sense as we previously discussed that a spreading process reaches nodes with a higher degree faster and only later reach lower-degree nodes. As the population of the highest fitness strains gets biased towards high-degree nodes, the mutation events in those strains are more likely to occur at high-degree nodes.

This can be expressed in a new equation for the speed of adaptation, using ⟨*τ*_*j*_ ⟩, denoting the time to the next adaptive mutation after an adaptive mutation in node *j*, and *π*_*j*_, that denotes the relative frequency of adaptive mutation events in node *j*:

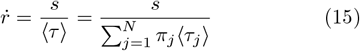

with 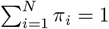.

Changing from ⟨*τ*⟩ to a location-dependent ⟨*τ*_*j*_⟩ is conceptually not very difficult and does not require us to make major changes to our approach. We can simply exchange ⟨*Y* ⟩(*t*) with an origin-dependent equivalent: ⟨*Y* ⟩ (*t* | *j*), the mean total of all sites occupied by strain *y*, if that strain originated from site *j*:

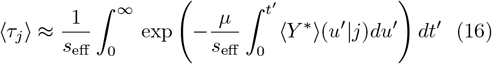

However, estimating *π* requires some additional considerations.

The approach we use here is a Markov process that models the transitions between adaptive mutation events and the sites of these events (as illustrated by Fig. 3). Because the waiting times between adaptive events are not necessarily exponentially distributed, the process is not a Markov chain but instead a more general Markov renewal process (Appendix E for the full definition).

**FIG. 3.**
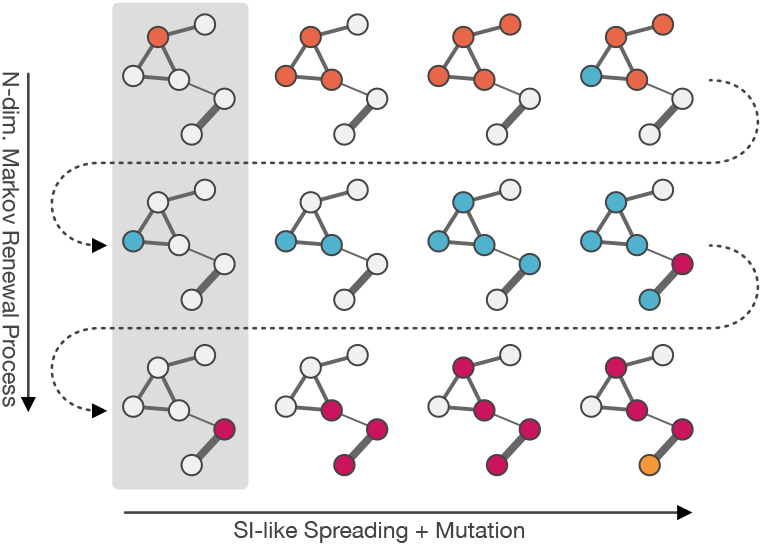
The dynamics of the emergence and spread of mutations through a heterogeneous network are complex. Our modeling approach: We focus on the emergence of a new strain with higher fitness than all others (adaptive mutation events). We construct a Markov renewal process modeling the jumps between these events and divide the dynamics into two parts: The 2^*N*^ -dimensional spreading dynamics (horizontal) and the *N* -dimensional transitions between origin sites of new adaptive mutations (vertical, highlighted in grey). Approximating the 2^*N*^ -dimensional spreading process using established methods from epidemic modeling yields the waiting times and transition probabilities between the *N* states of the Markov Renewal Process. See Section III C.

The process consists of a sequence of states, each state *S*_*i*_ corresponding to a possible origin node *i*. The waiting times between state transitions depend on the last state and are given by eq. 16.

The transitions probabilities between states are defined by the transition probability matrix *H*, where *H*_*ij*_ is the transition probability from state *S*_*j*_ to *S*_*i*_. From *H*, we compute the stationary distribution, *π*^*ζ*^, which models the distribution of the adaptive mutation events over all sites, by solving:

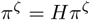

The probability vector *π*^*ζ*^ is an approximation of *π* and gives us the relative frequency of adaptive events in each site.

We define the transition probabilities using a similar approach to Eq. 4 and 6, but now we also consider the locations of mutation events:

We replace the mean number of all sites occupied by strain *y*, ⟨*Y*⟩ (*t*), with two explicitly location-dependent values: ⟨*Y* ⟩ (*t* | *j*) and ⟨*Y*_*i*_⟩ (*t* | *j*), the probability *of a single site i* being occupied by strain *y*, if strain *y* originated in site *j*. Note that 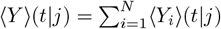.

Analogous to equations 4 and 14, the probability density of a *y* → *z* mutation event in site *i* following an *x* → *y* mutation event in site *j*, is then:

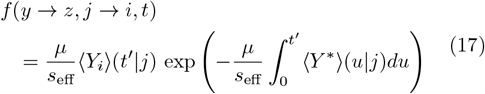

This gives us the equation:

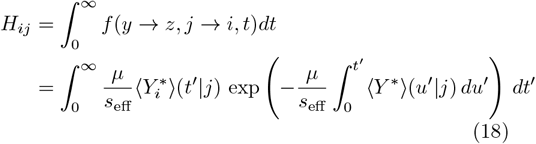

Note that in the equation of *H*_*ij*_ the biological parameters *µ* and *s*_eff_ only appear together as the fraction 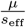.

For a given network, *H*_*ij*_ and by extension the distribution of adaptation mutations *π*^*ζ*^ depends only on a single biological parameter, the ratio between the mutation rate and the average fitness advantage of the fittest strain over its neighbors.

We can again estimate the values for 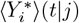 and 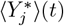 using approximation methods from network epidemiology, such as the heterogeneous mean-field (HMF) approximation.

#### 1. The relationship between origin frequency π_j_ and waiting time ⟨τ_j_ ⟩

This model explains an interesting approximate relationship between the frequency of adaptive mutations originating from a node, *π*_*j*_, and the mean waiting time to the next adaptive mutation, ⟨*τ*_*j*_⟩: *π*_*j*_ increases approximately linearly with the inverse of ⟨*τ*_*j*_⟩ (see Fig. 2d):

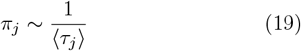

We explain this approximate relationship by comparing the Markov renewal process with a similar Markov chain process, for which this relationship holds exactly. The core idea is that each step in the Markov renewal process represents a “race”: The site *i* that both acquires the mutation from site *j* and and subsequently mutates “wins” and becomes the origin of the next adaptive mutation. Instead of the distributions of arrival time 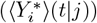 in equation 18, and 16, the Markov chain process will only include the mean waiting time of the entire event 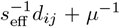, and model each competing event using a Poisson process. Here, the distance 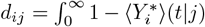 is the normalized mean arrival time to site *i* of the strain *y* with origin *j*.

The rate matrix *Q*^*C*^ characterizes the process:

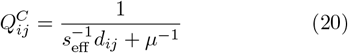

for off-diagonal elements and 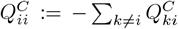. The arrival times *d*_*ij*_ = *d*_*ij*_ for SI dynamics are symmetric under the assumption of a symmetric network, *A*_*ij*_ = *A*_*ji*_ (see section II). Thus 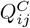 is symmetric and the Markov chain process fulfills detailed balance criteria. Based on these attributes, we can derive the following relationship between 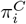 and 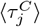:

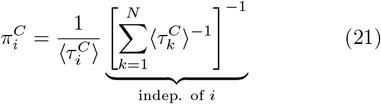

See Appendix E 1 for a detailed definition of the Markov chain process and the derivation of this equation.

Remarkably, this straightforward relationship carries over to the original model: the relationship between *π*_*i*_ and ⟨*τ*_*i*_⟩ ^−1^ remains approximately linear, albeit with a significant intercept, even for mutations rates with significant CI (see. Fig. 2c).

### D. The origin of the fittest strains depends on the time-frame given by the mutation rate

From Eq. 21, we can learn that the most likely origins of an adaptive mutation are biased towards origins that facilitate the spread of the mutation (as *π*_*i*_ ∝ ⟨ *τ*_*i*_ ⟩^−1^). Crucially however, the mutation rate (relative to *s*_eff_) dictates which parts of the function ⟨ *Y* ^*^⟩ (*t* | *j*) actually matter.

In some networks, the same nodes are either good or bad facilitators of spreading over all time scales. However, for many networks, this is not the case. A quintessential example of such a network is the dumbbell-like network. This network has two cores (highly connected subnetworks), that are themselves linked by a chain of nodes (see Fig. 4a). Nodes in the cores have many neighbors but reach the opposite core only slowly, while nodes in the middle of the chain have few neighbors but the smallest average distance to all others.

**FIG. 4.**
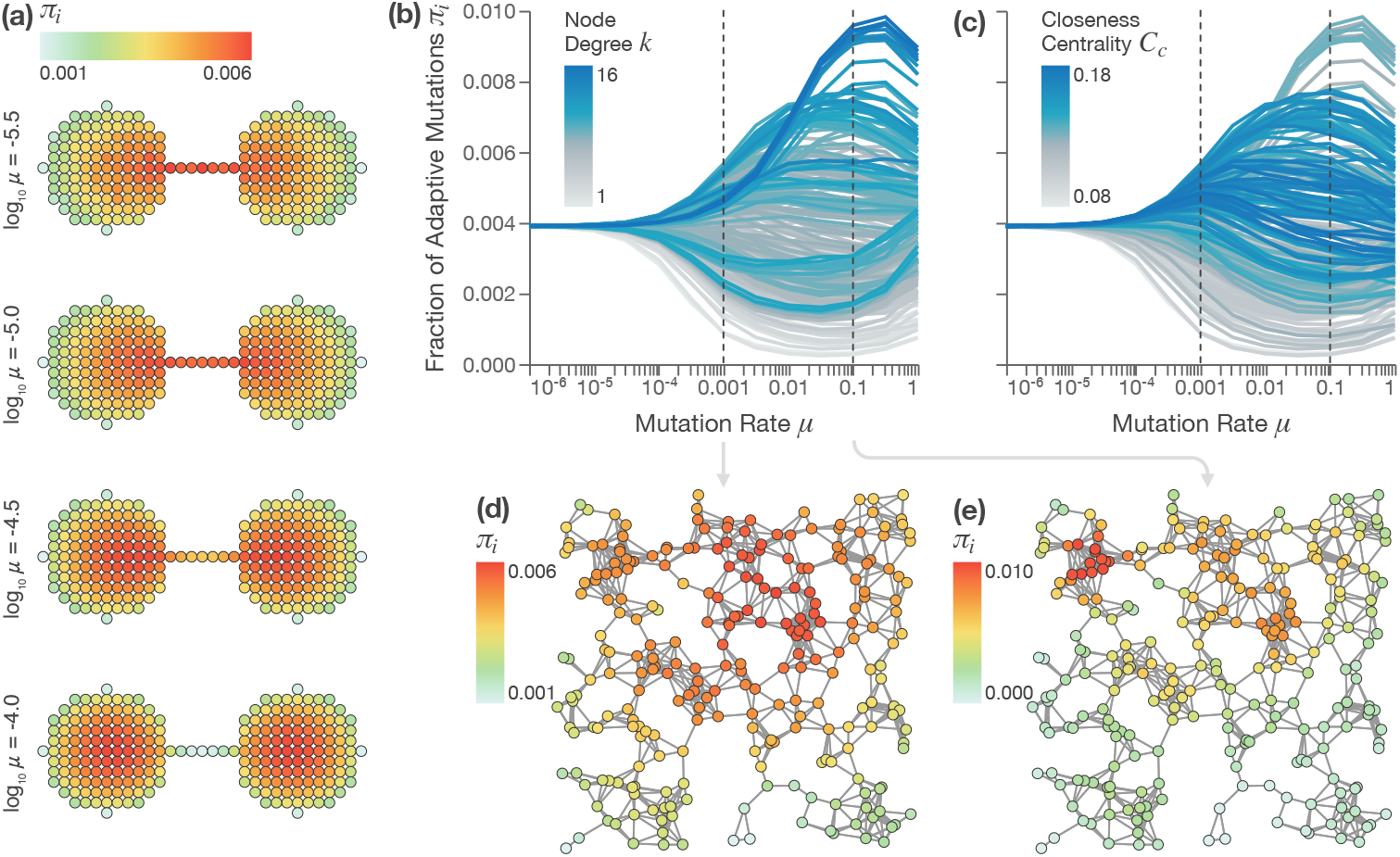
The origin of high-fitness strains shows a complex dependence on the mutation rate *µ* in different habitat networks. The mutations that produce mutants that have the highest fitness in the entire population, here called adaptive mutations, are biased towards nodes that facilitate the fast spread of the mutation. **(a)**: The distribution of adaptive mutations *π* for different mutation rates *µ* in a dumbbell-like network of two (grid-like) cores connected by a narrow bridge. This structure creates a mismatch of the local and global properties: Nodes located in the bridge are globally central but have few immediate neighbors, while nodes in the cores have many neighbors but are globally peripheral. For small *µ*, within the CI regime, global spreading properties dominate *π*: The nodes in the bridge are the most likely sources of adaptive strains. For larger *µ*, local spreading properties become more important and the distribution of adaptive mutations shifts to the center of the cores. **(b-e)**: This property is demonstrated for a habitat network generated via the Random Geometric Graph (RGG) model. **(b, c)**: The fraction of adaptive mutations *π*_*i*_ for all nodes *i* in relation to the mutation rate *µ*. The trajectories are highlighted depending on node degree *k*_*i*_ **(b)** or closeness centrality *C*_*c*_ **(c)**. This highlights how *π* generally correlates with *C*_*c*_ for low *µ* and with *k* for high *µ*. The black vertical dotted lines mark the values of *µ* shown in (d, e). **(d, e)**: Snapshots (at *µ* = 0.001 and *µ* = 0.1) of the distributions of adaptive mutations *π* in the RGG network. **(d)**: The regime with *µ* = 0.001 where a central cluster of nodes is the most likely origin. **(e)**: The regime with *µ* = 0.1, where the most likely origin is a peripheral cluster of a few highly connected nodes. Simulation results represent averages over 50,000 trajectories with a simulation time of *T* = 5*/µ, s* = 0.1 and *A*_*ij*_ = *a* = 1.

This means that ⟨*Y* ^*^⟩ (*t* | *j*) grows faster in the beginning for core node origins, but *y* fixates the fastest for chain nodes. In Fig. 4a, we demonstrate the consequences for the distribution of adaptive mutations *π*_*j*_. For low mutation rates *µ*, the most likely origin of adaptive mutations is the center of the bridge. As *µ* increases, the *π* distribution moves towards the centers of the core and for high *µ*, the middle nodes of the bridge are among the least likely origins of a new adaptive mutation.

A similar effect can be observed in a simple network model: the random geometric graph (RGG). In a RGG, nodes are embedded in Euclidean space and an edge is present when the inter-node distance is below a fixed threshold. The RGG, like the lattice, is one of the simplest spatial network models, but its randomness creates modularity and community structures.

Fig. 4, panels (b)–(e), explore the *π*-distribution for a RGG realization with 256 nodes. Similarly to the dumb-bell network, the distribution becomes more heterogeneous with increasing *µ*, but also qualitatively changes: For large *µ*, the node degree *k* is more indicative of high *π*_*i*_ values (Fig. 4b). For low *µ* the ordering of *π*_*i*_ closely follows the closeness centrality *C*_*c*_, the reciprocal of the sum of the shortest-path distances from the node to all others and a measure of global centrality (panel c).

Some approximation methods for ⟨*Y* ^*^ ⟩ (*t* | *j*) are inherently unable to capture these properties. In particular, approximation methods like the HMF approximation, used in section III A, reduce nodes to a micro-scale attribute, the node degree, and are therefore inherently unable to capture meso- or macroscopic spreading properties. However, other methods to approximate ⟨*Y* ^*^⟩(*t*|*j*) can capture these.

One is the dynamic message-passing algorithm (DMP), a state-of-the-art method to approximate epidemic dynamics [34, 52]. DMP tracks spreading probabilities along directed edges to incorporate direct-neighbor correlations but neglects higher-order dependencies that create the computational intractability of the exact 2^*N*^ - dimensional Markov Chain Process. It is provably exact on tree graphs and remains highly accurate on most sparse or networks with moderately few loops. Full equations are given in Appendix I.

The DMP approximation of *π* captures the changes with *µ* both qualitatively and quantitatively, as shown in Fig. 5 across multiple realizations of the RGG network (Fig. A5 for other realizations). For BA-type networks, the match is closer (Fig. A6). While the DMP successfully captures key aspects of CI across different regimes, it requires relatively costly numerical simulations. However, we can arrive at a much simpler, closed-form equation for ⟨*τ*_*j*_⟩ and *H*_*ij*_ by using a network distance based approximation, where each node pair is assigned a delay 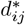, that captures the mean arrival time of a SI process from *j* to *i* (described in Appendix H 2). The results are nearly identical to the distribution-based approach for low *µ*, but deviates more strongly for larger *µ* (see Fig. 5b and Fig. A5).

**FIG. 5.**
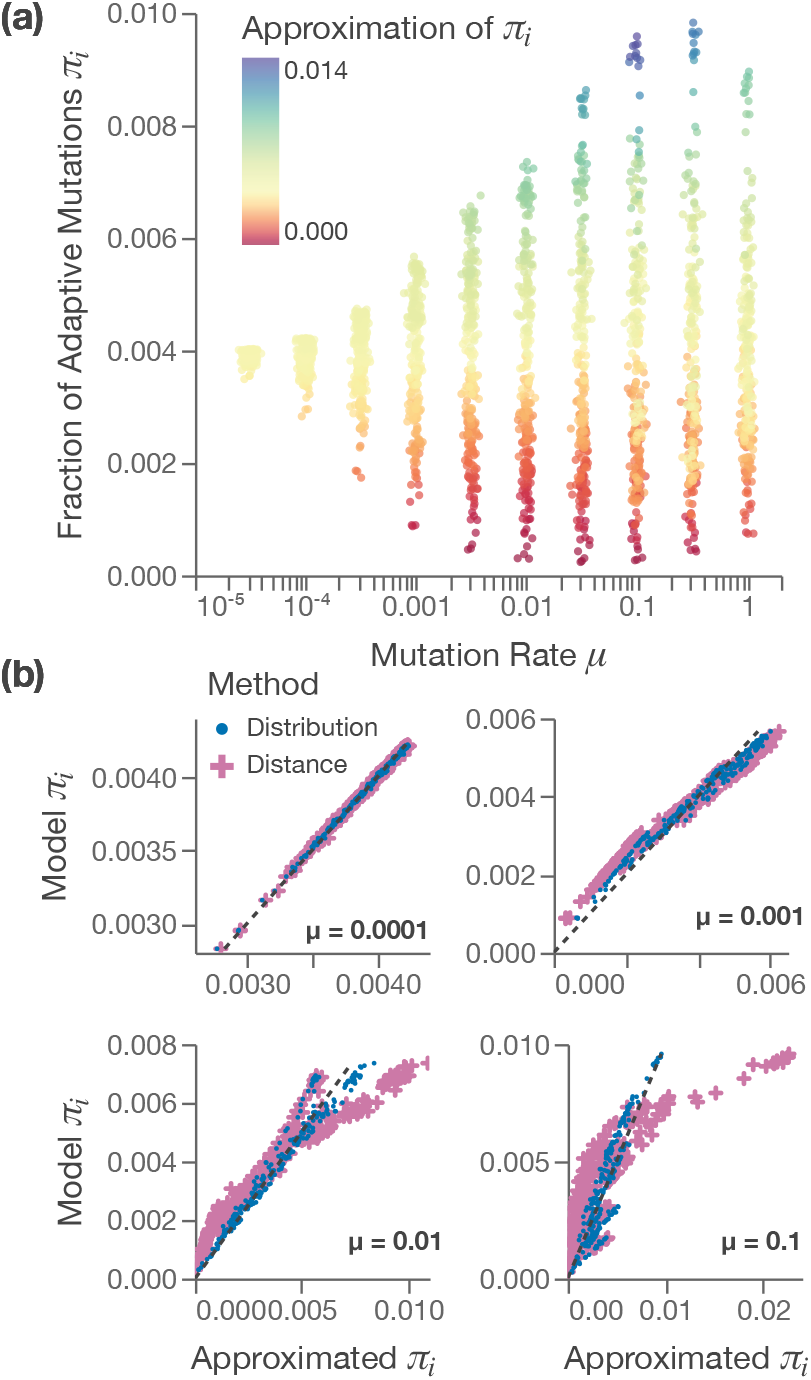
Estimates of the fraction of adaptive mutations for each node, *π*_*i*_, obtained via the Markov Renewal Process match the sampled results over a large range of *µ*. For large *µ* the heterogeneity is slightly overestimated (see (c)). As shown in (b), A distance-based approach, where the spreading dynamics are approximated by a single distance between each pair of nodes, performs well for small *µ* The network is the RGG with *N* = 256 from Fig. 4. Spreading approximated via dynamics message passing with average fitness difference, *s*_eff_ = *s*. Simulations represent averages over 50,000 trajectories with a simulation time of *T* = 5*/µ, s* = 0.1 and *A*_*ij*_ = *a* = 1. . Data points in (a) are displayed with a small random jitter along the x-axis to aid readability.

#### Effectiveness of control strategies under varying mutation rates

In the previous section we have demonstrated clearly that sites do not contribute equally to the speed of adaptation 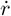 in the clonal-interference regime. Consequently, network modifications that exclude sites vary in their ef-fect on 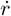, depending on how often the excluded sites produce high-fitness strains. Since the contributions of sites can vary with the mutation rate *µ*, the effects of network modifications will also vary with *µ*.

We use an illustrative example network, specifically constructed to contain multiple subnetworks that can be the most likely origins of the fittest strains, depending on the mutation rate *µ*. The network consists of multiple cores (fully-connected subnetworks) that are inter-connected by much weaker links (see inset in Fig. 6a). The central core is connected to three peripheral chains. The first chain consists of a single core, the second of two cores and the third of three. Within all cores, nodes are linked with *A*_*ij*_ = 1. All nodes in adjacent cores are linked with *A*_*ij*_ = 10^−4^, except for the two cores in the second chain, which are more strongly linked with *a* = 10^−2^. The single core in the first chain has 8 nodes, all others have 5.

**FIG. 6.**
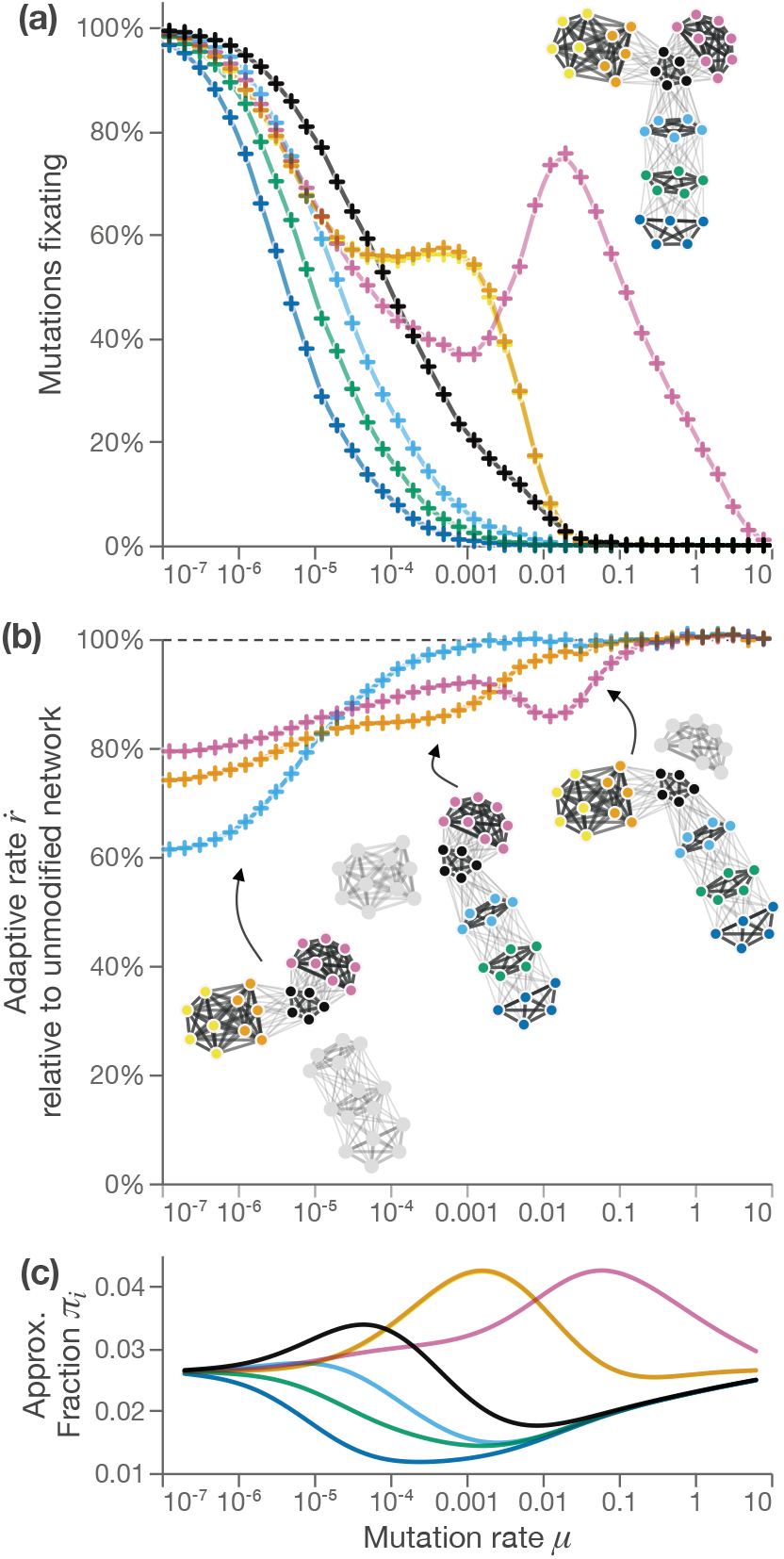
Multiple (*>* 2) distinct regimes and relative changes in adaptation speed 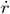 after network modifications and its dependence on mutation rate. The small example network is specifically designed so that it has three regions that optimize spreading for different timescales (see insert network representation **(a)**). **(a)**: Proportion of new strains fixating at various mutation rates *µ* for specific network regions. The colors represent the strain’s origin. **(b)**: Changes in the adaptive rate 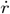 in the same network after three different modifications. Changes are shown relative to the 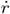 in the unmodified network. Each modification cuts either all links between the blue core (blue marks), the yellow cluster (yellow marks) or the pink cluster (pink marks) and the black core. Each modification has, for a certain range of *µ*, the strongest impact on 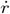. **(c)**: The approximated *π*_*i*_ values for the network replicate the different regimes observed in (a) and (b). All panels represent averages from 10,000 trajectories with a simulation time of *T/µ* = 5 and *s* = 0.1. Approximation via the DMP method on the network aggregated at core level.

We examine the effects of very specific modifications: the removal of each of the chains from the network, resulting in three different network versions. The networks and the effect of each modification on the adaptation speed 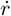 is illustrated in Fig. 6.

For all these modified structures, in the *strong-selection weak-mutation regime*, the decrease in the parameter 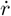 is directly proportional to the fraction of nodes that are no longer connected to the largest component. The modifications that remove more nodes are therefore more effective in reducing 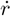. As *µ* increases, the difference between the modified and original networks decreases, as CI reduces the contribution of additional nodes to 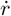 (as it would in every system with CI). Within the CI regime, we find that the influence of excluded sites on 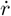 mirrors their contribution to fixating strains (see Fig. 6a for reference). Each of the three modifications has a regime in which it has the strongest effect on 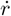. These regimes can be easily identified using our approximation of *π* (see Fig. 6c). The approximation uses the DMP method on the network aggregated at the core-level to avoid errors caused by the circular wiring within the cores.

## IV. CONCLUSIONS

Using a simple model of clonal interference (CI), we studied the dynamics of adaptation in spatially heterogeneous asexual populations. Our findings underscore the importance of network attributes, such as degree heterogeneity and average distances, for the speed of adaptation. They also highlight the interplay between the functional spreading attributes of the network across timescales and the biological parameters of the evolutionary process: The fittest lineages will be more likely to originate from a node or sub-network that optimizes the spread of mutations. However, for different mutation rates, the spreading dynamics on different timescales will determine which subnetworks are optimal. Small muta-tion rates will favor globally central origin nodes, while larger mutation rates will put a greater focus on shorter timescales and meso-or micro-scale node attributes like the degree.

These findings are supported by our proposed theoretical approach that models mutation events and their locations in the current fittest strain of the population with a Markov renewal process. We show how the transition probabilities and waiting times of this process are approximated using methods borrowed from epidemiology. This allows us to reduce a very complex system to one with a small state space of *N*, capturing the heterogeneity in the origin of what we call adaptive mutations—mutations that produce a strain with a higher fitness than previously present in the population. Because our approach works for arbitrary weighted networks, it can accommodate a large variety of population structures including meta-population structures.

While theoretically elaborate, our method can be simplified to systems that use distances to represent the arrival times of mutations to other sites, resulting in much simpler formulas and revealing an important and intuitive property: the likelihood of any site to produce an adaptive mutation, *π*_*i*_, is approximately inversely proportional to the waiting time for the next adaptive mutation, ⟨*τ*_*i*_⟩ (see section III C 1). This allows for the comparison of individual sites without solving the entire Markov renewal process by focusing on a single representative value, ⟨*τ*_*i*_⟩ (see section III C 1).

These findings have significant implications for interventions informed by evolutionary dynamics. The effectiveness of modifications to the network that hinder or facilitate adaptation can vary greatly depending on the mutation rate. This has potential applications in understanding and manipulating the evolutionary trajectories of asexual populations in fields such as epidemiology and conservation biology. For example, understanding the effects of network modifications on adaptation speed could inform strategies for managing ecosystem resilience or controlling the spread of pathogens where clonal interference phenomena and the origin of new variants are relevant [53–56]. Especially in the case of multi-year pandemics, seasonal changes in case numbers and gradual changes in infection rates could change the balance between mutation rate and spreading rates over time and lead to shifts in optimal intervention strategies.

A key limitation of our method is the assumption of a fixed fitness increment introduced by each mutation. While an exponential distribution of fitness increments is generally regarded as more realistic [3], the observed patterns change only marginally with exponentially distributed fitness increments (see Appendix F 1), likely because an exponential fitness distribution still results in a clearly defined timescale for the spreading of new mutations. Expanding our methodology to variable fitness distributions could help clarify this, with a key part of such an expansion being the inclusion of more than two fitness levels, which would also improve the accuracy of the existing method for higher mutation rates. This could be achieved through a Markov Renewal Process that models transitions between sequences of origins, with states being pairs of sites and the transition probability matrix *H*_*k*←*j,j*←*i*_ describing the likelihood that a transition between site *i* to site *j* is followed by a transition from site *j* to site *k*.

Future work could examine more complex and realistic population structures, investigating whether the trade-off between local and global centrality that creates multiple distinct regimes in our example networks is a common real-world network property. Prior research has shown that the world-wide air transportation network indeed features a separation of the most central and most connected cities that could cause such a trade-off [57].

Additionally, experimental validation of our theoretical predictions would be invaluable for testing the applicability of our model to real-world populations. Theoretically proposed clonal interference phenomena have been experimentally observed in viral, bacterial, and eukaryote cultures [7, 58–63]. One relatively simple experimental system could involve bacteria growing in a dumbbell-like culture, i.e., a structure with two large dishes connected by a thin bridge of substrate, where sites on the bridge offer increased global centrality but a lower number of close neighbors. Alternatively, a similar setup to [7] could be used, where different meta-population topologies were created by manually transferring samples between isolated cultures.

In conclusion, our study unveils a complex interplay between network heterogeneity, mutation rate, and clonal interference in asexual populations. We demonstrate that the mutation rate is a key factor in determining the relative fixation rates of strains across different network regions. Based on the arrival times of mutations between network sites, we developed a comprehensive theoretical framework enabling us to analyze and predict these dynamics on arbitrary network structures. This work opens new avenues for understanding and manipulating the evolution of asexual populations, with far-reaching implications spanning from ecosystem management to epidemic control.

## APPENDIX A Network Construction

In this appendix we provide full specifications for each network model used in our comparisons in section III A.

All networks were normalized so that the average weighted degree

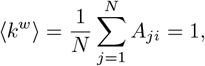

and, for ER, BA, and grid networks, the average *unweighted* degree (i.e. the average number of neighbors) was fixed at ⟨*k*⟩ = 8.

### Erdős–Rényi (ER) networks

There are two variations of the ER network: In one, each possible link is randomly present or absent with a fixed probability. In the second, the number of links, *M*, is fixed and each link is randomly assigned to a set of nodes, without duplicates. To make comparisons to other population structures easier, we use the second definition and avoid any variability in the overall rate of invasion events. Additionally, we discard networks with more than one component. That means there is always a path between any node-pair.

### Barabási–Albert (BA) networks

The BA network is generated by a growth model, where the network is generated in steps, each adding nodes and/or links. In case of the BA model, each step adds exactly one node and a fixed number of links (*m*) that connect the new node to existing ones. Key is that new links are biased to nodes with already a high number of links: the preferential attachment rule. This rule generates a scale-free link-distribution and mimics a key mechanism in the real world: For example, a new member of a social network is more likely to establish social links to community members that are socially active and already well-connected [44].

Barabási–Albert graphs are grown by sequentially adding nodes with (m) edges each, attached preferentially to high-degree nodes. To compare fairly across topologies, we enforce an exact edge count (*M* = 4*N*, ⟨*k*⟩ = 8) for the BA instances, which a standard BA((N,4)) from the default star seed would miss by a constant offset (*M* = 4*N* − 16). We therefore adopt a two-stage construction that preserves linear preferential attachment: First, we build a connected seed (*G*_0_) with *N* = 16 and *m* = 8, which has 64 edges. Then, we continue BA growth to *N* with *m* = 4 using *G*_0_ as the initial graph. This choice slightly strengthens the earliest hubs but leaves the degree tail and ultra-small-world behavior intact and will have little impact on large graphs. Implementation using the NetworkX python package [64].

### Planar 8-neighbor grid

The planar grid configuration is a planar 8-neighbor grid (aka King’s grid, as each link is a legal move of a king chess piece). The grid has periodic boundary conditions, so each node is truly identical.

### Well-mixed population

In the well-mixed model every pair of nodes is directly linked. Our normalization of 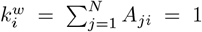 is equivalent to maintaining a constant encounter density as *N* grows: In the terms of the well-mixed model of individuals that occupy some volume and randomly encounter each other, the volume increases proportional with *N*, so that the density stays the same. This normalization corresponds to the usual chosen parametrization of a well-mixed system, where the initial growth rate is independent of the population size *N* and fixation in the absence of competitors occurs on a timescale ∝ log *N* .

## Appendix B Gillespie’s Stochastic Simulation Algorithm and its Application to Our Model

Gillespie’s stochastic-simulation algorithm (SSA) is an exact procedure for sampling the time evolution of any well-mixed Markov process in which every elementary reaction is a Poisson process. It was originally developed to simulate chemical reactions and has been widely adopted in a variety of fields including computational systems biology and epidemiology. Its logic rests on the following assumptions:

1. **Markov property:** each elementary process is memoryless and free of explicit time-dependence
2. **Independence:** all processes are statistically independent of one another
3. **Uniqueness of events:** in an infinitesimal interval *dt* at most one event can occur.

When these conditions hold, two key results follow:

1. At any time *t* the waiting time Δ*t* until the next event of *any* type is exponentially distributed with the mean of Λ^−1^, where Λ is the total sum of all rates.
2. The probability of any specific event (*i*) to be the next event is exactly *λ*_(*i*)_*/*Λ, where *λ*_(*i*)_ is the rate of the Poisson process associated with the event (*i*).

Both of these steps can be easily implemented, and by repeating the draw-time / choose-event cycle, one obtains a statistically exact trajectory that respects the continuous-time dynamics without introducing an artificial time step. Note that the rates might change after each cycle and might have to be recomputed.

In our model, elementary events fall into two classes: (a) invasions *X*_*v,i*_ + *X*_*u,j*_ → *X*_*u,i*_ + *X*_*u,j*_ occur at rate *A*_*ij*_(*r*_*u*_ − *r*_*v*_); (b) mutations at node *i* occur at rate *µ* and increase fitness by *s* (see Section II for the full definition). Applying the SSA to the model in Section II, where elementary events fall into these two classes, we follow the four stages illustrated in Fig. A1.

**FIG. A1.**
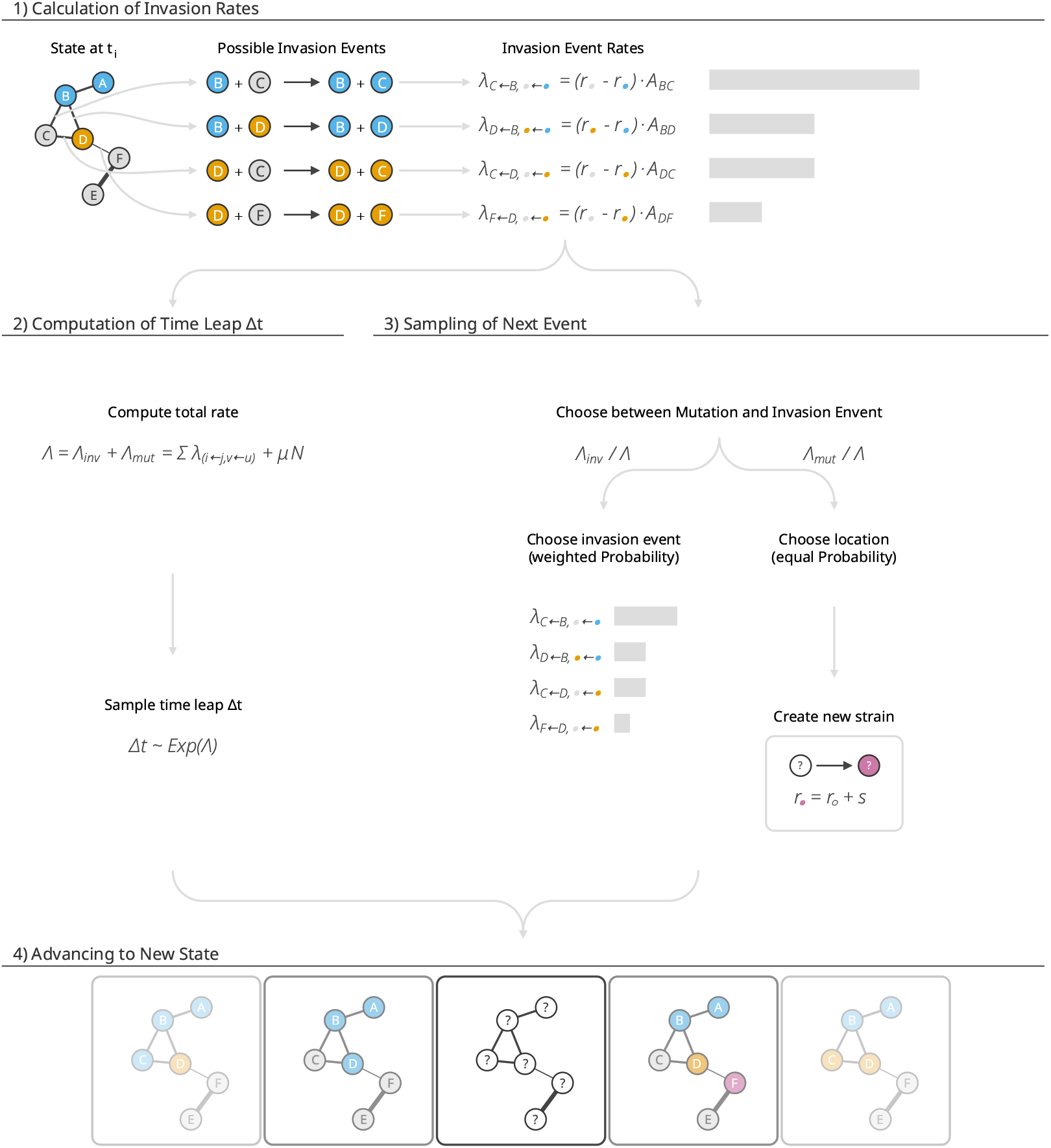
Gillespie algorithm for coupled mutation–invasion dynamics on a weighted network. (1) Current configuration n(nodes colored by resident strain) with the set of feasible invasion events and their rates *λ*_*i*←*j,v*←*u*_. Note that each event always corresponds to a link between nodes occupied by strains of different fitness values *r*. (2) The combined event rate Λ = Λ_inv_ + Λ_mut_ defines an exponential waiting-time distribution from which the time leap Δ*t* is drawn. (3) The next event is chosen by first deciding between mutation (probability Λ_mut_*/*Λ) and invasion (probability Λ_inv_*/*Λ). Conditional on an invasion, one of the candidate invasion events is selected with probability proportional to its rate. For a mutation event, a node is chosen uniformly and a new strain with fitness increment *s* is created. (4) The network and simulation time *t* is updated and the procedure then repeats from panel (a).

### 1. Rate calculation

Every node mutates at the constant rate *µ*, giving a time-independent total mutation rate Λ_mut_ = *µN* . Thus, we don’t need to update the mutation rates in our base model. Possible invasions are attached to directed links *j* → *i* for which the donor strain *u* is fitter than the recipient strain *v*. Each carries the rate

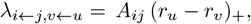

where *A*_*ij*_ is the (symmetric) link weight and (·)_+_ is the positive-part function that is zero for negative arguments. Summing these rates yields the invasion flux Λ_inv_(*t*). A crucial efficiency point is that, because only one node changes in any event, only the links incident to that node acquire new rates, and after the first cycle we only need to iterate over these links for updates.

### 2. Time leap

The waiting time to the next event is drawn from

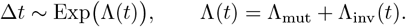

### 3. Choosing the event

With probability Λ_mut_*/*Λ the next change is a mutation and with the complementary probability Λ_inv_*/*Λ it is an invasion. A mutation is resolved by picking a node *i* uniformly at random and replacing its resident strain *u* by a novel strain *w* with fitness *r*_*w*_ = *r*_*u*_ + *s*. For an invasion, a link is selected with probability *λ*_*i*←*j,v*←*u*_*/*Λ_inv_. The fitter strain *u* then replaces *v* at node *i*.

### 4. State and time update

The global clock is advanced to *t* ← *t* + Δ*t* and the affected node is updated to the new state and the cycle restarts at stage (a).

### 5. Modified implementations

Within this paper we use variations of our model that generally represent simplifications of the original.

#### a. Double Mutation Simulation

The first one is a variant model that only considers three different clones (called *x, y* and *z*) with different fitness values: *r*_*z*_ = *r*_*y*_ + *s* = *r*_*x*_ + 2*s* (referenced in section III B).

We start these simulations from an initial state that has a single node occupied by clone *y* and all others occupied by *x*. The model does not include *x* → *y* mutation events, but the clone *y* can both spread by invasion events and mutate in a *y* → *z* mutation event. After the first *y* → *z* mutation event, the simulations are stopped.

Note that because this variant does not include *x* → *y* mutations, the rate Λ_mut_ will not be constant, but will have to be updated after every step with Λ_mut_ = *µY* (*t*). Additionally, nodes still in state *x* are excluded when sampling the location of a mutation event. Otherwise, the simulations are identical.

#### b. Fittest-Clone Mutation Simulation

This model is not limited to a predefined set of clones, but also restricts mutation events: Only the fittest clone in the population can mutate. This means that the models behavior is closer to our approximation that similarly neglects the influence of mutations in other clones. Again, the rate Λ_mut_ will not be constant, but will have to be updated after every step and the nodes not occupied by the fittest clone are excluded when sampling the location of a mutation event.

## Appendix C Notation and symbols

**TABLE 1.**
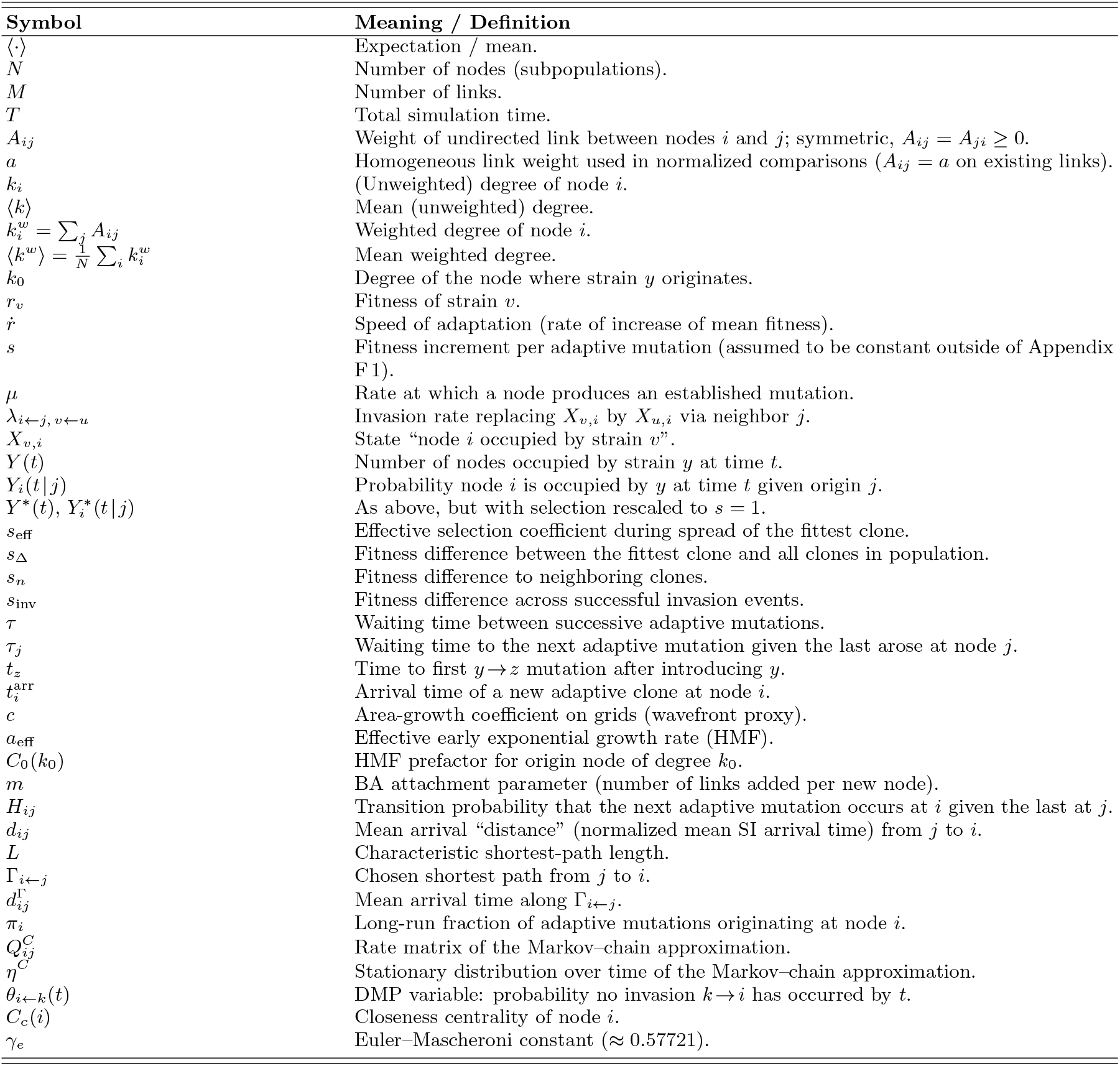
Symbols used throughout the paper. Rates are per unit time.

## Appendix D Approximations

### 1. The influence of *x* → *y* Mutations on *t*_*z*_ and 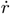

Within our theoretical considerations we neglect the influence of mutations on the growth of ⟨*Y*⟩ (*t*).

This approximation is similar to the approach taken in previous research [10, 35] and assumes that the nonlinear growth by invasion/competition will generally far outweigh the linear rate of growth by mutations.

**TABLE 2.** Acronyms used in the text.

|  |  |
| --- | --- |
| CI | Clonal interference |
| ER | Erdős-Rényi network |
| BA | Barabási-Albert network |
| RGG | Random geometric graph |
| WM | Well-mixed population |
| SSA | Gillespie's stochastic simulation algorithm |
| HMF | Heterogeneous (degree-based) mean-field |
| DMP | Dynamic message passing |
| MRP | Markov renewal process |

Starting from a homogeneous population (⟨*t*_*z*_ ⟩in section III B), this assumption will quickly lead to large deviations (see Fig. A2(a)). However, in the full model with a distribution of different fitness values, this assumption becomes more accurate, as there are fewer clones that lag behind the fittest clone only by a single fitness increment *s* (see Fig. A2(b)).

**FIG. A2.**
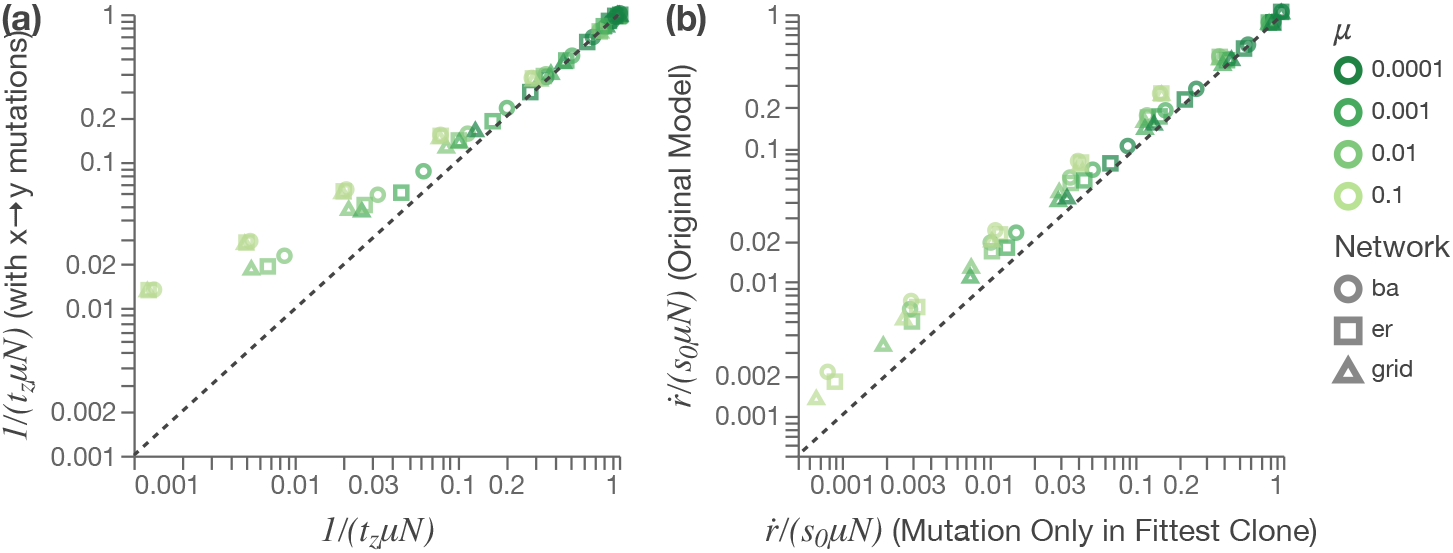
This figure shows the influence of mutations outside of the fittest clone in the system. In panel (a) we focus on a system with three clones of increasing fitness, *x* → *y* → *z*. Starting with a homogeneous population of *x* and a single clone of *y*, we are interested in how quickly the first *y* → *z* mutation occurs (see quantity ⟨*t*_*z*_ ⟩ in section III B and section B 5 a), and more specifically, the slowdown compared to the sequential accumulation of mutations, 1*/*(*t*_*z*_⟩ *µN*), where *µN* is the total rate of mutation events. In one scenario, *y* will only spread by invading nodes occupied by clone *x* (x-axis), in the other, nodes can also transition from *x* → *y* via mutation events (y-axis). Shown are 3 networks (BA, ER, and Grid, see section III A and A), 5 sizes (*N* = [16, 64, 256, 1024, 4096]) and 4 mutation rates (*µ* = [10^−4^, 0.001, 0.01, 0.1]). For values close to 1 (little slowdown), ⟨*t*_*z*_⟩ will not deviate much between the scenarios. For values with a stronger slowdown (1*/*(⟨*t*_*z*_⟩ *µN*) ≪ 1), the deviations become more pronounced, particularly for the grid networks. In panel (b) we show the slowdown of the rate of adaptation (compared to the speed of sequential accumulation, *sµN*) in the full model (y-axis) and a modified version without mutations outside the fittest strains (x-axis, see appendix section B 5 b). Shown are the same 3 networks (BA, ER, and Grid, see section III A and A), 5 sizes (*N* = [16, 64, 256, 1024, 4096]) and 4 mutation rates (*µ* = [10^−4^, 0.001, 0.01, 0.1]). Comparatively, the differences between the models are smaller.

### 2. Mean-field Approximation of *Y* (*t*) and its influence on ⟨*t*_*z*_ ⟩

In our derivation of equation 7, we used a mean-field approximation and approximated the mean of the integral with the integral of the mean value of *Y* (*t*):

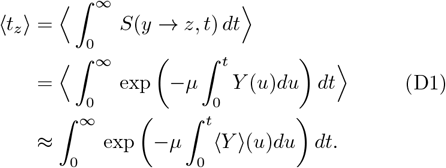

We show the validity of this approximation for 3 network models and different *µ* in Fig. A3. The deviations between the simulated values ⟨*t*_*z*_⟩ and the approximated values using equation 7 are generally small but we see some deviations from the simulated results, particularly for very small *µ* in large heterogeneous networks.

**FIG. A3.**
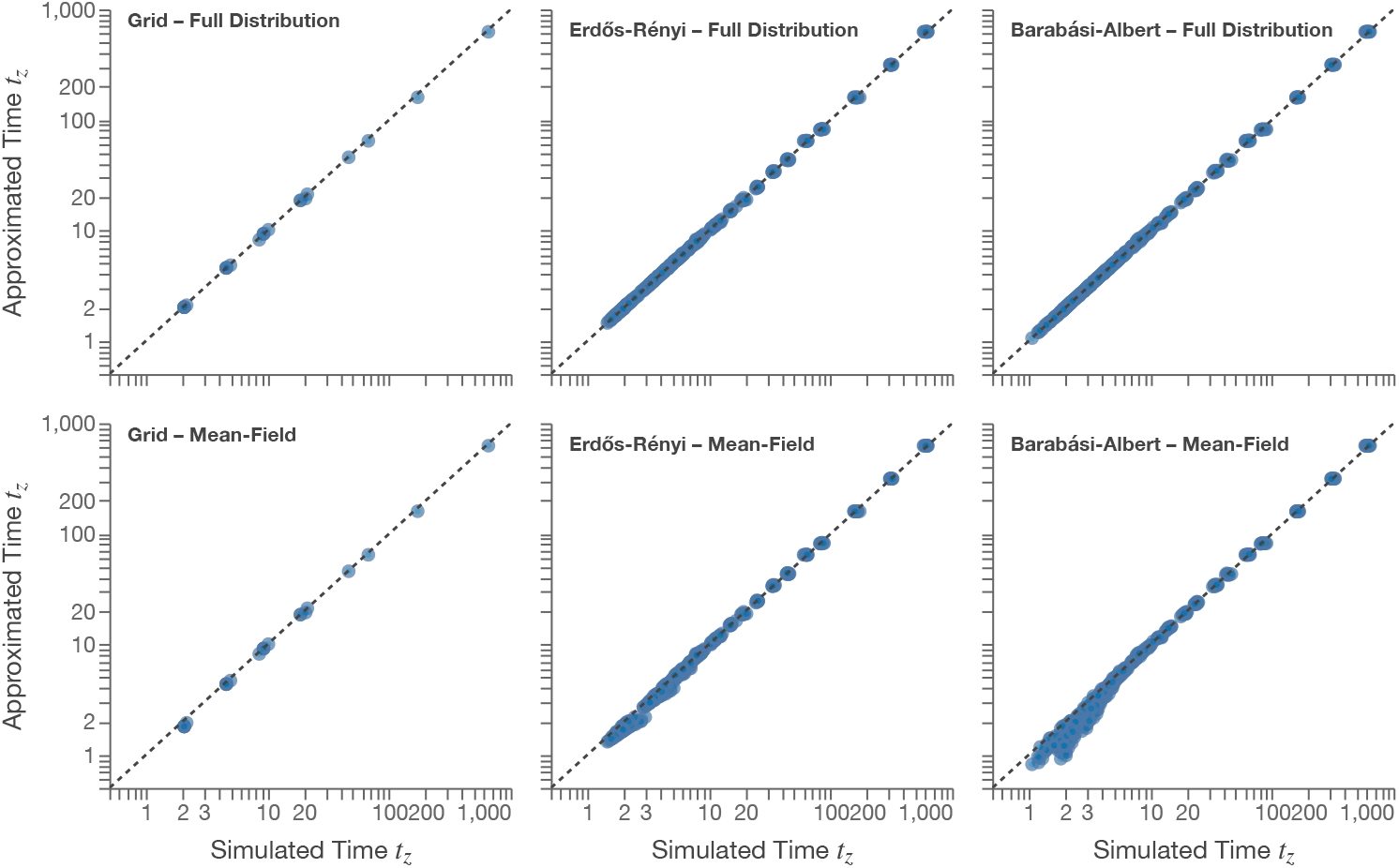
This figure shows the deviation between simulated and approximated ⟨*t*_*z*_⟩ values for 3 different network models. The upper row shows the result of equation 6, with no systematic deviation from the simulated results as this equation is describing the simulation results exactly. The lower row shows the result of equation 7 where a mean-field approximation was used to describe the variable *Y* (*t*). For the ER and BA network, ⟨ *t*_*z*_⟩ values for *µ* = [10^−4^, 10^−3^, 10^−2^, 10^−1^] and *N* = [2^4^, 2^5^, 2^6^, …, 2^14^] and from 10 different starting locations are shown. For the grid network, only *N* values that are perfect squares (*N* = [2^4^, 2^6^, 2^8^, …, 2^14^]) were used and the simulations were not repeated for different origins (as they would be identical in nature). The values of *Y* (*t*) were generated via sampling (1,000 trajectories) to isolate the contribution of the mean-field approximation. The simulated values of ⟨*t*_*z*_ ⟩ likewise represent averages over 1,000 trajectories.

### 3.2 Effect of *k*_0_ on the expected value of ⟨ *t*_*z*_⟩^*ER*^

Starting from Eq. 11 with 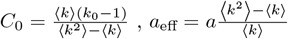 and ⟨*k*^2^⟩= ⟨*k*^2^⟩+ ⟨*k*⟩:

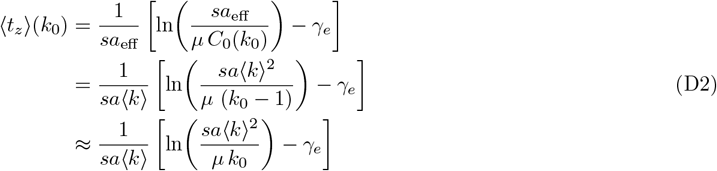

so that, after marginalizing over *k*_0_,

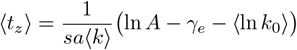

Assume *k*_0_ is drawn from a zero-truncated binomial Bin(*N, p*), mean ⟨*k*⟩ = *Np* (based on the ER network model). A standard delta-method expansion of ln *k*_0_ around ⟨*k*⟩ yields

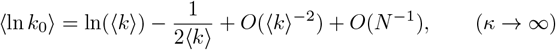

so replacing ⟨ln *k*_0_⟩ by ln⟨*k*⟩ in the expression for ⟨*t*_*z*_⟩ in Eq. 12 produces an error of

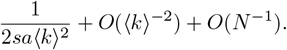

Because the correction is ≈ 1*/*(2*s*⟨*k*⟩^2^) and the Gumbel scale is ≈ 1*/*(2*s*⟨*k*⟩), the relative error in units of the intrinsic scale is ≈ 1*/*(2⟨*k*⟩), which is typically negligible once ⟨*k*⟩ is moderate (⟨*k*⟩ *>* 10).

## Appendix E Markov Renewal Process for Origins of Adaptive Mutations

This appendix provides the definition of the Markov renewal process used in Section III C to model the sequence of *adaptive* mutation events and their locations.

The Markov process *ζ*_*t*≥0_(*t*) models the transitions between adaptive mutation events and their corresponding sites.

The process consists of a sequence of states, [*S*_1_, …, *S*_*n*_], and corresponding jump times [*T*_1_, …, *T*_*n*_]. The state space of *ζ*_*t*≥0_(*t*), 𝒮 = {*S*_*i*_ : *i* = 1, 2, 3, …, *N* }, has *N* elements, each element *S*_*i*_ corresponding to an adaptive mutation event at node *i*. The jump times are associated with the waiting times 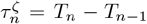 between state transitions. The process is memoryless: The next state and the waiting times depend only on the current state.

The process is defined by the following probability density of the next adaptive event occurring at node *i* at time *t* following the last event at *j*:

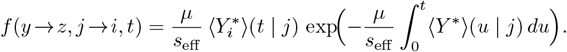

Integrating over *t* yields the transition probabilities

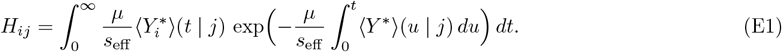

where *H*_*ij*_ denotes the transition probability from state *S*_*j*_ to *S*_*i*_. The mean waiting time for leaving state *S*_*j*_ is given by

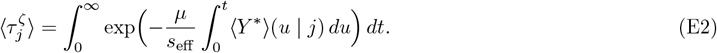

### 1. Markov chain process 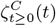 and the relationship between 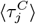, and *π*^*C*^

By replacing the full waiting-time distributions of the renewal process with their mean transition rates, we derive a corresponding continuous-time Markov chain (see section III C 1).

We denote the Markov Chain Process approximation of *ζ*_*t*≥0_(*t*) with 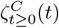. The process 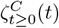 shares the same state space 𝒮 and is characterized by the rate matrix:

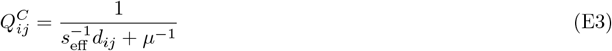

As the arrival times *d*_*ij*_ = *d*_*ji*_ for SI dynamics are symmetrical (under the assumption of a symmetrical network, *A*_*ij*_ = *A*_*ji*_, see section II), *Q*^*C*^ is also symmetrical. Equivalent entities for 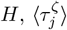, and *π*^*ζ*^ are denoted 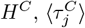, and *π*^*C*^, respectively.

Equations 18 and 16 simplify to:

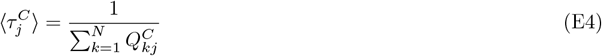

and

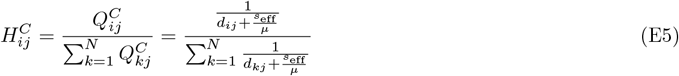

In a Markov chain process, at the steady state, the rate at which the process enters any state *i* can be expressed as 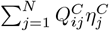 and it must match the rate at which the process exits 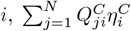, where *η*^*C*^ is the stationary distribution of 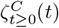 (*t*) *over time* (as opposed to the stationary distribution *over adaptation events, π*^*C*^). As *Q*_*ij*_ is symmetric, the process 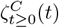 fulfills detailed balance criteria and the homogeneous distribution 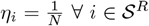 is the trivial solution for the steady-state *η*^*C*^.

The distribution *π*_*i*_ can be expressed as the rate 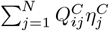 normalized over all events:

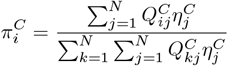

with 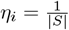 and 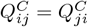 and equation E4:

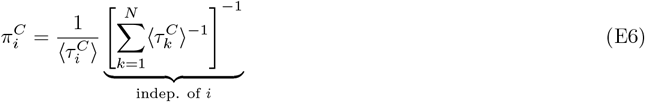

Hence, 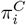 (the probability that we enter the state *S*_*i*_ compared to any other state) is determined entirely by the mean waiting times 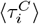 (as it is the inverse of the rate at which we exit the state *S*_*i*_). In this simplified system, the mean waiting times are entirely sufficient to understand the model’s broader behavior.

## Appendix F Additional Simulation Results

### 1. Exponentially distributed fitness increment *s* ∼ Exp

If we replace the constant fitness increment *s* = *s* with an increment drawn from an exponential distribution, *s* ∼ exp(⟨*s*⟩), the broad patterns are very similar (see Fig. A4). The distribution of *π* also has very similar dependencies on the mutation rate *µ*, with global centrality dominating for small *µ* and a shift to nodes with strong local properties for large *µ*. Overall we see a slight decrease in heterogeneity and the transition to local properties shifts to higher mutation rates *µ*.

**FIG. A4.**
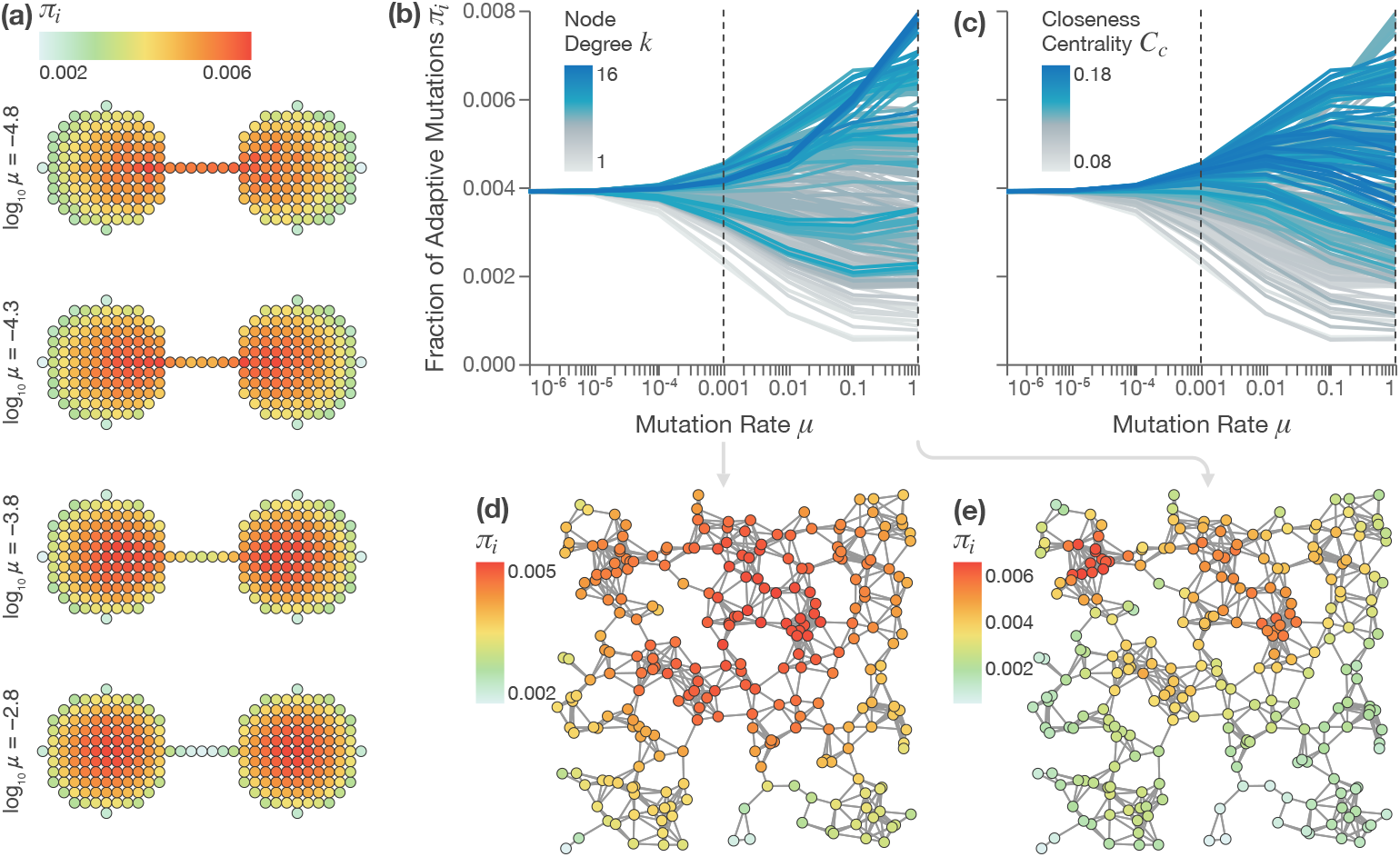
This figure is a reproduction of Fig. 4 of the main article using an exponentially-distributed fitness increment *s* ∼ Exp with a rate parameter of 0.1. Again, the origin of high-fitness strains shows a complex dependence on the mutation rate *µ* in different habitat networks. The mutations that produce mutants that have the highest fitness in the entire population, here called adaptive mutations, are biased towards nodes that facilitate the fast spread of the mutation. **(a)**: The distribution of adaptive mutations *π* for different mutation rates *µ* in a dumbbell-like network of two (grid-like) cores connected by a narrow bridge. This structure creates a mismatch of the local and global properties: Nodes located in the bridge are globally central but have few immediate neighbors, while nodes in the cores have many neighbors but are not globally central. For small *µ*, within the CI regime, global spreading properties dominate *π*: The nodes in the bridge are most likely source of adaptive strains. For larger *µ*, local spreading properties become more important and the distribution of adaptive mutations shifts to the center of the cores. **(b-e)**: This property is demonstrated for a habitat network generated via the Random Geometric Graph (RGG) model. **(b, c)**: The fraction of adaptive mutations *π*_*i*_ for all nodes *i*, in relation to the mutation rate *µ*. The trajectories are highlighted depending on node degree *k*_*i*_ **(b)** or closeness centrality *C*_*c*_ **(c)**. This highlights how *π* generally correlates with *C*_*c*_ for low *µ* and with *k* for high *µ*. The black vertical dotted lines mark the values of *µ* shown in (d, e). **(d, e)**: Snapshots (at *µ* = 0.001 and *µ* = 0.1) of the distributions of adaptive mutations *π* in the RGG network. **(d)**: The regime with *µ* = 0.001 where a central cluster of nodes is the most likely origin. **(e)**: The regime with *µ* = 0.1, where the most likely origin is a peripheral cluster of a few highly linked nodes. Simulation results represent averages over 50,000 trajectories with a simulation time of *T* = 10*/µ*.

### 2. Repetition of Approximation results for other seeds

For RGG (Fig. A5) and BA (Fig. A6), the Markov–renewal + DMP approximation closely tracks sampled *π*_*i*_ over a wide range of *µ* (with *s*_eff_ = *s*) across independent seeds for the network construction. The distance–based variant agrees well at small *µ* but degrades at large *µ* (hence omitted for *µ* = 0.1). Agreement is generally closer on BA than on RGG.

**FIG. A5.**
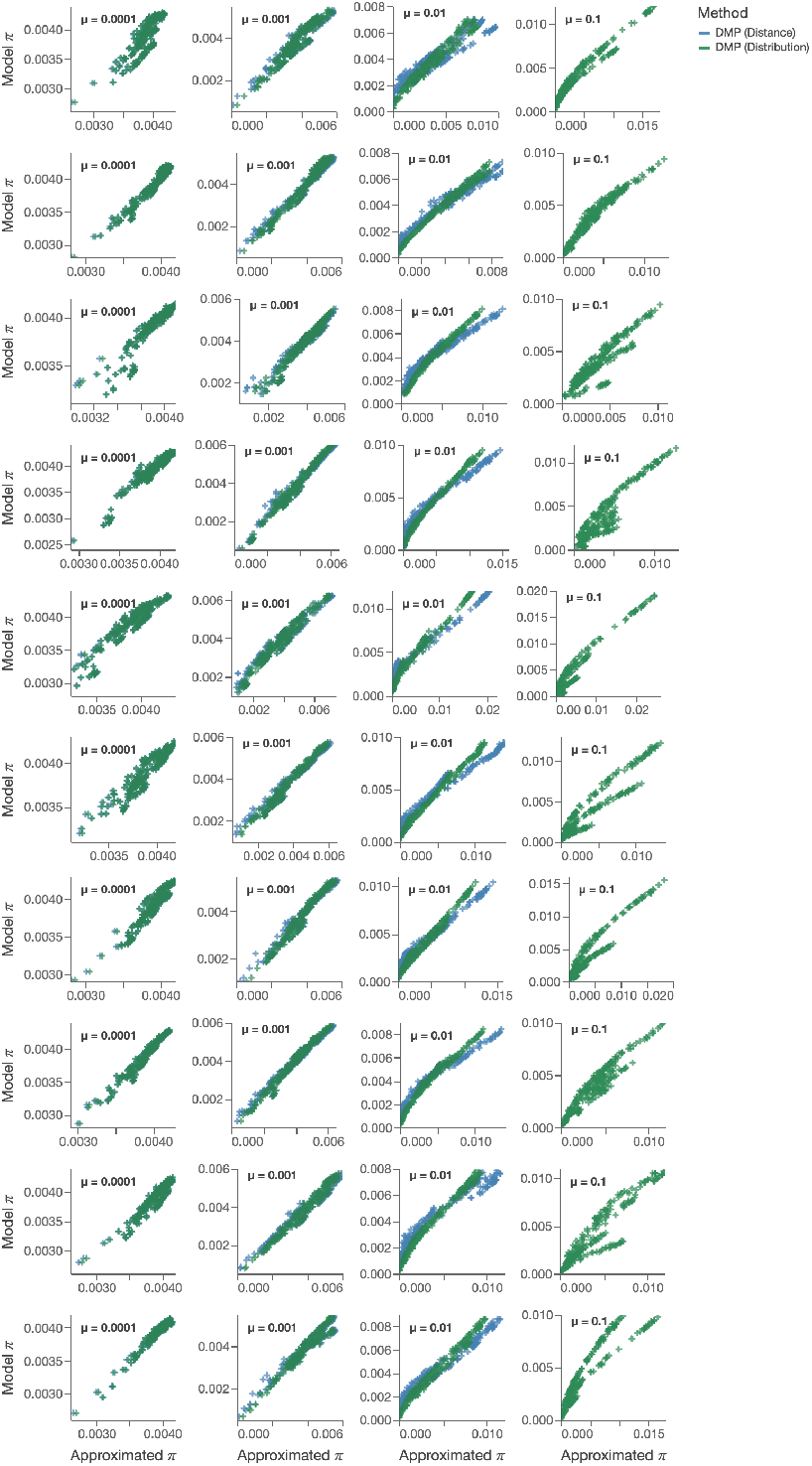
Estimates of the fraction of adaptive mutations for each node, *π*_*i*_, obtained via the Markov Renewal Process match the sampled results over a large range of *µ*. Each row represents a different realization of the RGG network with the same parameters as the network from Fig. 4. Blue marks show the results of the distance based approximation (omitted for *µ* = 0.1). Spreading approximated via dynamics message passing with average fitness difference, *s*_eff_ = *s*. Simulations represent averages over 50,000 trajectories with a simulation time of *T* = 5*/µ*.

**FIG. A6.**
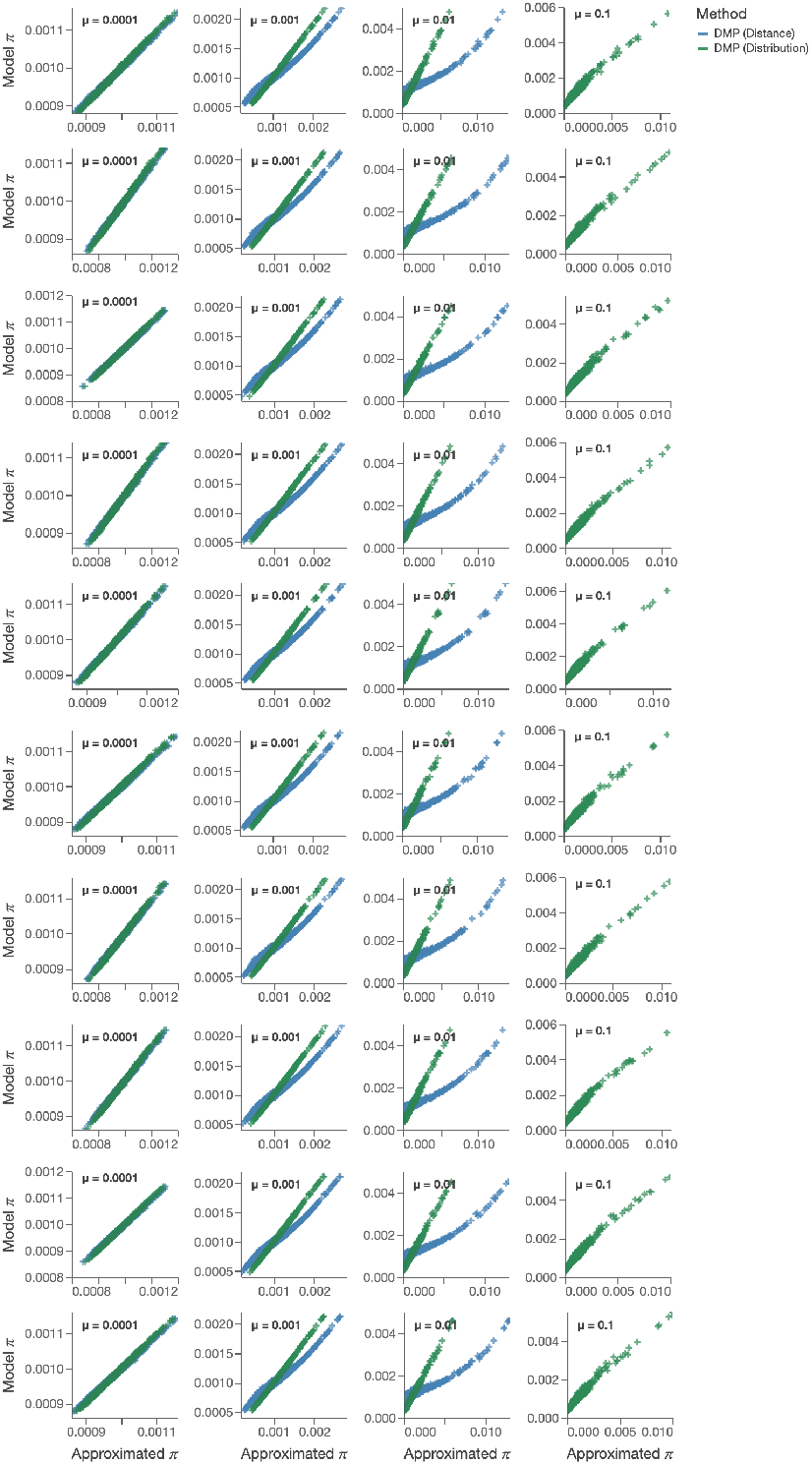
Estimates of the fraction of adaptive mutations for each node, *π*_*i*_, obtained via the Markov Renewal Process match the sampled results over a large range of *µ*. Each row represents a different realization of the BA network with the same parameters as the network from Fig. 3. Blue marks show the results of the distance based approximation (omitted for *µ* = 0.1). Spreading approximated via dynamics message passing with average fitness difference, *s*_eff_ = *s*. Simulations represent averages over 50,000 trajectories with a simulation time of *T* = 5*/µ*.

## Appendix G Estimating *s*_eff_ via Balance of Adaptation and Invasion Rates

In the full model, a new mutation does not spread in a homogeneous background but rather in a population containing potentially many different clones, each with distinct fitness values. While the exact distribution of encountered fitness values is difficult to determine, one can approximate the effect by replacing the baseline selection coefficient *s* with an effective value *s*_eff_ (see Figure A7, which uses sampled values for *s*_eff_).

**FIG. A7.**
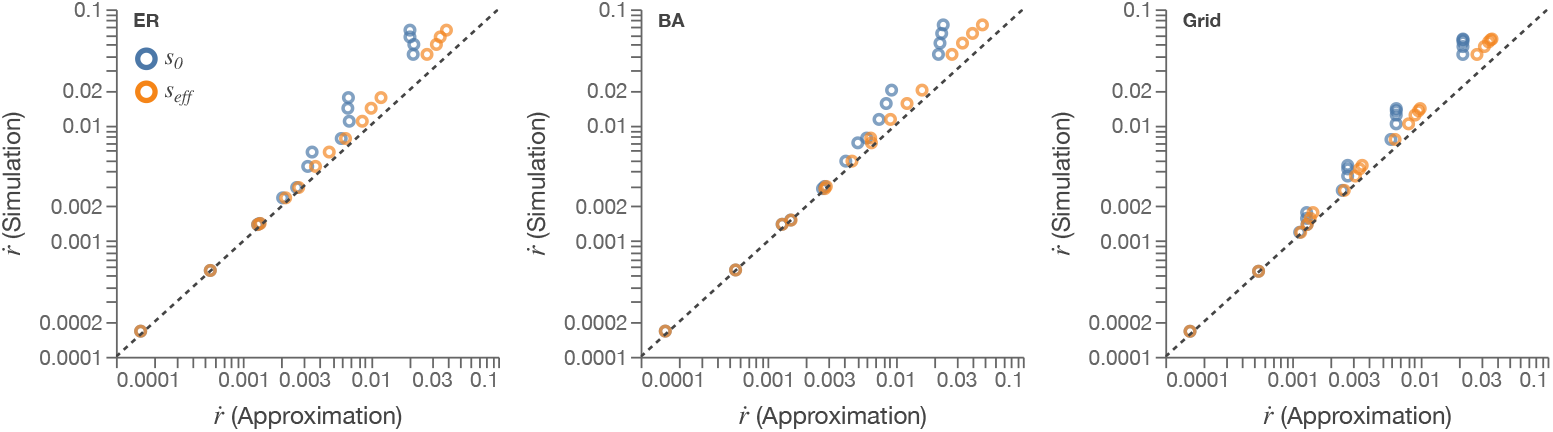
Comparison of estimated values of the speed of adaptation 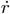 (x-axis) to simulated values of 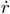 (y-axis) in networks generated by three different network models. Each estimated value was computed via Eq. 16 either with *s*_eff_ set to *s* = 0.1 or to a sampled *s*_eff_ and sampled values for ⟨*Y* ^*^⟩ (*t*). The sampled values of *s*_eff_ were generated by choosing random time points and then choosing uniformly from the links between a node occupied by a clone of maximum fitness and any lower-fitness neighbor, if such a link exists. Using sampled values of *s*_eff_ increases the accuracy of predictions compared to using *s*_eff_ = *s*.

To estimate *s*_eff_ without simulating the model, one can use the balance between the adaptation rate based on the fittest clones in the population, Eq. 16, and the increase in mean fitness in the whole population, as shown in [10, 46] for well-mixed populations. When *µ* ≪ *sA*_*ij*_, the increase in mean fitness in the whole population will be mainly driven by invasion events and will be affected by the increasing variance in fitness differences in three ways:

1. New mutations spread faster, since they encounter more nodes occupied by strains with a large fitness difference to the spreading strain.
2. The fitness difference between invading and resident clones will be larger on average, and the fitness increase per invasion event will increase.
3. New mutations spread, on average, only to a subset of nodes, biased towards those reached faster.

On the other hand, the rate of adaptation 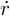 via adaptive mutation is less strongly affected, because the speed-up only comes from the faster spreading of the adaptive mutations. We can therefore estimate *s*_eff_ by finding the value of *s*_eff_ for which:

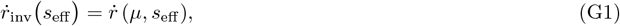

where 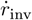 denotes the average fitness increase in the population via invasions alone.

We present a heuristic here for 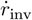, contingent on a number of approximations that allow us to estimate *s*_eff_ in populations that are close to the well-mixed case in the sense that the average path distances are small and path degeneracy is high.

To develop this heuristic, we must track several relevant distributions of fitness differences and approximate their relationships. We denote the distribution of fitness differences between the spreading strain and all other clones as *s*_Δ_. The distribution of fitness differences between the spreading strain and all neighboring clones is denoted as *s*_*n*_. This is generally smaller than *s*_Δ_. Finally, the distribution of fitness differences between the spreading strain and all neighboring clones that are invaded is denoted as *s*_inv_.

Several key relationships govern these distributions. First, note that *s*_inv_ *> s*_*n*_ since lower fitness neighbors are more likely to be invaded. Indeed, the distribution is exactly *f* (*s*_inv_) = *s*_*n*_*f* (*s*_*n*_) and ⟨*s*_inv_⟩ = ⟨*s*_*n*_⟩ + Var(*s*_*n*_)*/* ⟨ *s*_*n*_⟩ . Second, since fitness differences can only accumulate if invasion events are less frequent per node than adaptive mutations, an adaptive mutation will, on average, reach only *Ns/*⟨*s*_inv_⟩ nodes. Third, 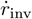 will scale linearly with ⟨*s*_inv_⟩, while both adaptation rates will scale with ⟨*s*_*n*_⟩ through *Y* (*t*).

We make the following simplifications. First, we assume that *s*_Δ_ decreases at a constant rate of 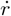. Second, we assume that the fitness differences to the highest fitness clone are approximately Poisson distributed. From that follows ⟨*s*_inv_⟩ ≈ ⟨*s*_*n*_⟩ + *s*. Third, we assume that *s*_*n*_ ≈ *s*_Δ_, based on the approximation of high path degeneracy.

For a given value of *s*_eff_, the average fitness difference ⟨*s*_inv_⟩ is approximately:

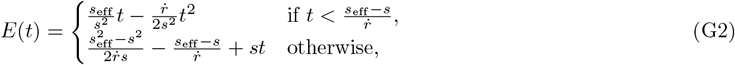

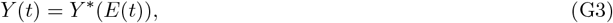

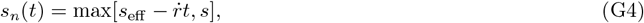

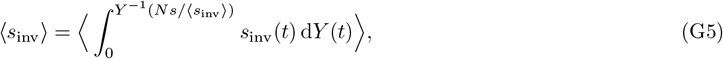

where *Y* ^−1^(·) is the inverse function of *Y* (·).

By numerically solving these equations together with Eq. G1, we can estimate *s*_eff_ (see Fig. A8).

**FIG. A8.**
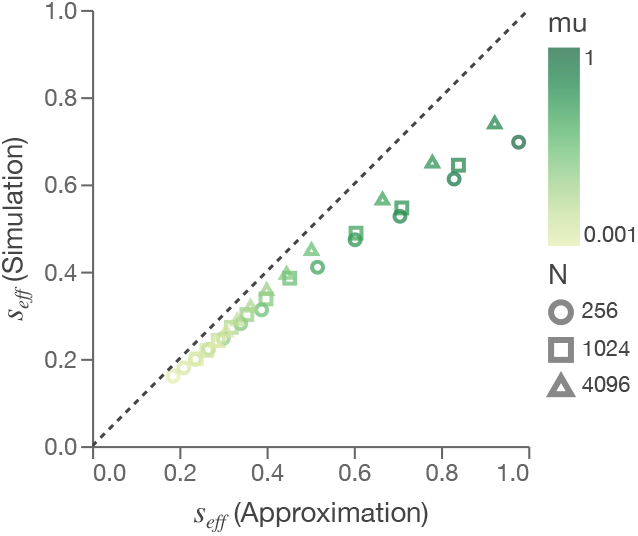
Comparison of estimates of *s*_eff_ (x-axis) to simulated values of *s*_eff_ (y-axis) in BA networks. Estimates were computed via the heuristic detailed in Appendix G.

## Appendix H Estimation of *Y* (*t*)

### 1. Shortest-Path Based Approximation of the arrival time distribution

The rates along the shortest path between two nodes can be used as a simple approximation of the arrival time of a spreading process in a network.

We define the ordered path Γ = { *A*_(1)_, …, *A*_(*L*)_ } with elements *A*_*ℓ*_ being the elements *A*_*ij*_ of the adjacency matrix along the path.

We choose the path that minimizes the path length 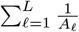 as the shortest path to *i* from *j*:

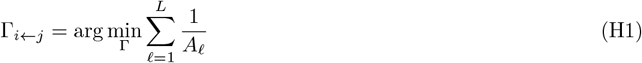

Let 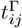 be the random variable describing the arrival time of a normalized SI-type process starting in *j* to *i* that travels only along the shortest path Γ_*i*←*j*_, with the spreading rate set to 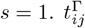 will be the sum of *L* exponentially distributed random variables: 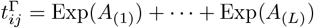, a hypoexponentially distributed variable.

The distribution of 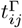 can be expressed as a special case of a phase-type distribution. Using ***α*** = [1, 0, …, 0], **1** is a column vector of ones of size *L*, and

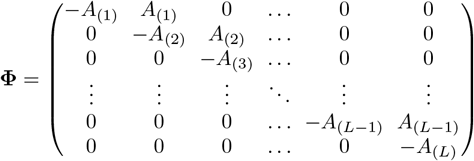

we can write the CDF of 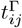 as

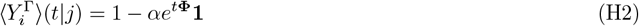

and the mean arrival time as

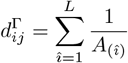

The distribution 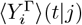 can then serve as an approximation of 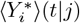, the CDF of the arrival time along any path.

### 2. Distance-based approximation of 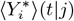 **and** ⟨*Y* ^*^⟩(*t*|*j*)

In this simplified formulation, we replace the true cumulative distribution function 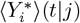 of the arrival time 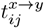 by a unit step at its mean 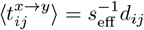. Concretely, we define

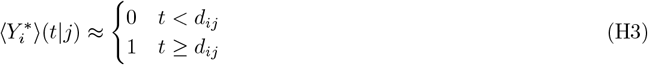

so that all stochastic variability is collapsed into a single deterministic jump at *d*_*ij*_.

Using this assumption, ⟨*Y* ^*^⟩ (*t* |*j*) can be expressed as piece-wise exponential decay, where we arrange the delays *d*_1*j*_, *d*_2*j*_, …, *d*_*Nj*_ in increasing order of magnitude such that:

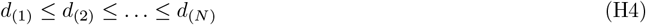

where *d*_(1)_ denotes the smallest delay in magnitude and *d*_(*N*)_ denotes the largest. Additionally we define *d*_(0)_ = 0, *d*_(*N*+1)_ = ∞, and *d*_(*m*)_ = *d*_*i*←*j*_. Now ⟨*τ*_*j*_⟩ and *H*_*ij*_ simplify to the following closed-form equations:

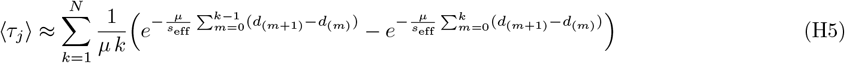

and

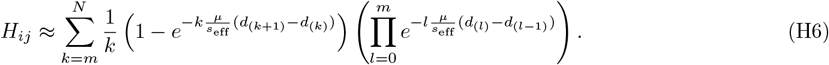

## Appendix I Dynamic Message-Passing (DMP) Equations for approximating 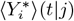 and ⟨*Y* ^*^⟩(*t*|*j*)

The equations in this Appendix are mathematically entirely equivalent to the DMP equations for a standard SI epidemic system (as described in [34]) but translated to the spreading of a strain *y* through a homogeneous population of a strain *x* based on the dynamics outlined in our model (in the absence of mutations) with a normalized fitness difference of *r*_*y*_ − *r*_*x*_ = 1.

We consider an undirected network of *N* nodes with nonnegative weights *A*_*ij*_ = *A*_*ji*_.

All nodes are either occupied by strain *x* or by the fitter strain *y* and the marginal probabilities of the two states are 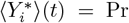 (node *i* is occupied by strain *y* at time *t*) and 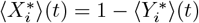.

For each directed edge *k* → *i*, we define

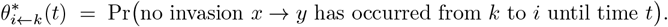

We also introduce the following two additional edge-based variables:

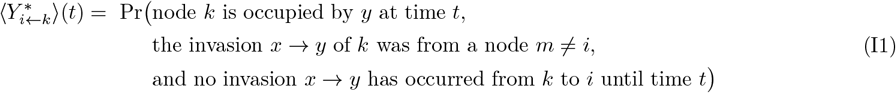

This expresses the potential of *k* to transmit the state *y* to *i* at time *t*.

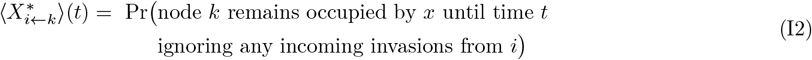

This is the most un-intuitive of the three edge-based variables but can be used to calculate 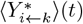 from 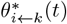 via:

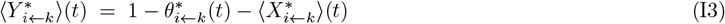

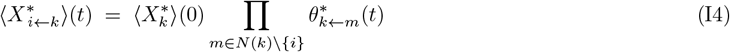

With the invasion rate *A*_*ik*_, the rate equation for *θ*_*i*←*k*_(*t*) is:

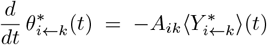

Together, these define the continuous-time DMP system. The node-based probabilities are given by:

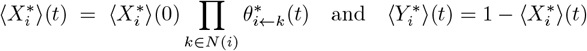

With 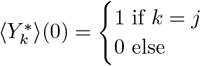 we arrive at the quantities 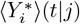 and 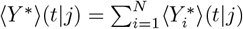.

## References

[1] D. M. McCandlish and A. Stoltzfus, Modeling evolution using the probability of fixation: history and implications, The Quarterly review of biology 89, 225 (2014).

[2] J. H. Gillespie, Some properties of finite populations experiencing strong selection and weak mutation, The American Naturalist 121, 691 (1983).

[3] P. J. Gerrish and R. E. Lenski, The fate of competing beneficial mutations in an asexual population, in Mutation and Evolution, Vol. 7, edited by R. C. Woodruff and J. N. Thompson (Springer Netherlands, Dordrecht, 1998) pp. 127–144, series Title: Contemporary Issues in Genetics and Evolution.

[4] P. D. Sniegowski and P. J. Gerrish, Beneficial mutations and the dynamics of adaptation in asexual populations, Philosophical Transactions of the Royal Society B: Biological Sciences 365, 1255 (2010).

[5] J. A. G. De Visser and D. E. Rozen, Clonal interference and the periodic selection of new beneficial mutations in escherichia coli, Genetics 172, 2093 (2006).

[6] K. C. Kao and G. Sherlock, Molecular characterization of clonal interference during adaptive evolution in asexual populations of saccharomyces cerevisiae, Nature genetics 40, 1499 (2008).

[7] P. P. Chakraborty, L. R. Nemzer, and R. Kassen, Experimental evidence that network topology can accelerate the spread of beneficial mutations, Evolution Letters 7, 447 (2023).

[8] P. R. A. Campos, P. S. C. A. Neto, V. M. de Oliveira, and Gordo, Environmental heterogeneity enhances clonal interference, Evolution 62, 1390 (2008).

[9] V. L. Chaves Filho, V. M. de Oliveira, and P. R. Campos, Adaptation of asexual populations in correlated environments, Physica A: Statistical Mechanics and its Applications 389, 5725 (2010).

[10] M. M. Desai, D. S. Fisher, and A. W. Murray, The speed of evolution and maintenance of variation in asexual populations, Current biology 17, 385 (2007).

[11] E. A. Martens and O. Hallatschek, Interfering Waves of Adaptation Promote Spatial Mixing, Genetics 189, 1045 (2011).

[12] E. Lieberman, C. Hauert, and M. A. Nowak, Evolutionary dynamics on graphs, Nature 433, 312 (2005).

[13] F. Alcalde Cuesta, P. González Sequeiros, and Á. Lozano Rojo, Suppressors of selection, PLoS One 12, e0180549 (2017).

[14] M. Broom, J. Rychtář, and B. T. Stadler, Evolutionary Dynamics on Graphs - the Effect of Graph Structure and Initial Placement on Mutant Spread, Journal of Statistical Theory and Practice 5, 369 (2011).

[15] C. J. Paley, S. N. Taraskin, and S. R. Elliott, Temporal and dimensional effects in evolutionary graph theory, Physical Review Letters 98, 098103 (2007).

[16] F. Fu, L. Wang, M. A. Nowak, and C. Hauert, Evolutionary dynamics on graphs: Efficient method for weak selection, Physical Review E 79, 046707 (2009).

[17] L. Marrec, I. Lamberti, and A.-F. Bitbol, Toward a Universal Model for Spatially Structured Populations, Physical Review Letters 127, 218102 (2021).

[18] M. Frean, P. B. Rainey, and A. Traulsen, The effect of population structure on the rate of evolution, Proceedings of the Royal Society B: Biological Sciences 280, 20130211 (2013).

[19] J. Tkadlec, A. Pavlogiannis, K. Chatterjee, and M. A. Nowak, Population structure determines the tradeoff between fixation probability and fixation time, Communications Biology 2, 138 (2019).

[20] J. Tkadlec, A. Pavlogiannis, K. Chatterjee, and M. A. Nowak, Fast and strong amplifiers of natural selection, Nature Communications 12, 4009 (2021).

[21] C. Pokalyuk, L. A. Mathew, D. Metzler, and P. Pfaffelhuber, Competing islands limit the rate of adaptation in structured populations, Theoretical Population Biology 90, 1 (2013).

[22] R. A. Fisher, On the dominance ratio, Proceedings of the royal society of Edinburgh 42, 321 (1923).

[23] R. Fisher, The evolution of dominance; reply to professor sewall wright, The American Naturalist 63, 553 (1929).

[24] S. Wright, Evolution in mendelian populations, Genetics 16, 97 (1931).

[25] S. Wright, Statistical genetics and evolution, Bull. Amer. Math. Soc. 48(4), 223 (1942).

[26] E. de A. Goncalves, V. M. de Oliveira, A. Rosas, and P. R. Campos, Speed of adaptation in structured populations, The European Physical Journal B 59, 127 (2007).

[27] L. Perfeito, I. Gordo, and P. R. Campos, The effect of spatial structure in adaptive evolution, The European Physical Journal B - Condensed Matter and Complex Systems 51, 301 (2006).

[28] C. Paley, S. Taraskin, and S. Elliott, The two-mutant problem: clonal interference in evolutionary graph theory, Journal of chemical biology 3, 189 (2010).

[29] J. Otwinowski and J. Krug, Clonal interference and muller’s ratchet in spatial habitats, Physical Biology 11, 056003 (2014).

[30] D. Brockmann and D. Helbing, The Hidden Geometry of Complex, Network-Driven Contagion Phenomena, Science 342, 1337 (2013).

[31] F. Iannelli, A. Koher, D. Brockmann, P. Hövel, and M. Sokolov, Effective distances for epidemics spreading on complex networks, Physical Review E 95, 012313 (2017).

[32] E. Cator and P. Van Mieghem, Second-order mean-field susceptible-infected-susceptible epidemic threshold, Phys. Rev. E 85, 056111 (2012).

[33] A. S. Mata and S. C. Ferreira, Pair quenched mean-field theory for the susceptible-infected-susceptible model on complex networks, Europhysics Letters 103, 48003 (2013).

[34] B. Karrer and M. E. Newman, Message passing approach for general epidemic models, Physical Review E—Statistical, Nonlinear, and Soft Matter Physics 82, 016101 (2010).

[35] M. De Domenico, Diffusion geometry unravels the emergence of functional clusters in collective phenomena, Physical review letters 118, 168301 (2017).

[36] C. Hens, U. Harush, S. Haber, R. Cohen, and B. Barzel, Spatiotemporal signal propagation in complex networks, Nature Physics 15, 403 (2019).

[37] G. Bertagnolli and M. De Domenico, Diffusion geometry of multiplex and interdependent systems, Physical Review E 103, 042301 (2021).

[38] M. Boguna, I. Bonamassa, M. De Domenico, S. Havlin, D. Krioukov, and M. Á. Serrano, Network geometry, Nature Reviews Physics 3, 114 (2021).

[39] J. B. S. Haldane, A mathematical theory of natural and artificial selection, part v: Selection and mutation, Mathematical Proceedings of the Cambridge Philosophical Society 23, 838–844 (1927).

[40] D. T. Gillespie, Exact stochastic simulation of coupled chemical reactions, The journal of physical chemistry 81, 2340 (1977).

[41] P. ERDdS and A. R&wi, On random graphs i, Publ. math. debrecen 6, 18 (1959).

[42] P. Erdos, A. Rényi, et al., On the evolution of random graphs, Publ. math. inst. hung. acad. sci 5, 17 (1960).

[43] E. N. Gilbert, Random graphs, The Annals of Mathematical Statistics 30, 1141 (1959).

[44] A.-L. Barabási and R. Albert, Emergence of scaling in random networks, Science 286, 509 (1999).

[45] R. Pastor-Satorras and A. Vespignani, Epidemic dynamics and endemic states in complex networks, Physical Review E 63, 066117 (2001).

[46] M. M. Desai and D. S. Fisher, Beneficial mutation– selection balance and the effect of linkage on positive selection, Genetics 176, 1759 (2007).

[47] M. Boguá, R. Pastor-Satorras, and A. Vespignani, Epidemic spreading in complex networks with degree correlations, in Statistical mechanics of complex networks (Springer, 2003) pp. 127–147.

[48] M. Barthelemy, A. Barrat, R. Pastor-Satorras, and A. Vespignani, Velocity and hierarchical spread of epidemic outbreaks in scale-free networks, Physical Review Letters 92, 178701 (2004).

[49] M. Barthélemy, A. Barrat, R. Pastor-Satorras, and A. Vespignani, Dynamical patterns of epidemic outbreaks in complex heterogeneous networks, Journal of theoretical biology 235, 275 (2005).

[50] D. Smilkov, C. A. Hidalgo, and L. Kocarev, Beyond network structure: How heterogeneous susceptibility modulates the spread of epidemics, Scientific reports 4, 4795 (2014).

[51] H. Yang, M. Tang, and T. Gross, Large epidemic thresholds emerge in heterogeneous networks of heterogeneous nodes, Scientific reports 5, 13122 (2015).

[52] A. Y. Lokhov, M. Mézard, and L. Zdeborová, Dynamic message-passing equations for models with unidirectional dynamics, Physical Review E 91, 012811 (2015).

[53] M. Amicone and I. Gordo, Molecular signatures of resource competition: Clonal interference favors ecological diversification and can lead to incipient speciation*, Evolution 75, 2641 (2021).

[54] B. H. Good, S. Martis, and O. Hallatschek, Adaptation limits ecological diversification and promotes ecological tinkering during the competition for substitutable resources, Proceedings of the National Academy of Sciences 115, E10407 (2018).

[55] N. Strelkowa and M. Lässig, Clonal Interference in the Evolution of Influenza, Genetics 192, 671 (2012).

[56] P. P. Klamser, V. d’Andrea, F. Di Lauro, A. Zachariae, S. Bontorin, A. Di Nardo, M. Hall, B. F. Maier, L. Ferretti, D. Brockmann, and M. De Domenico, Enhancing global preparedness during an ongoing pandemic from partial and noisy data, PNAS Nexus 2, pgad192 (2023), <https://academic.oup.com/pnasnexus/article-pdf/2/6/pgad192/51011871/pgad192.pdf>.

[57] R. Guimera, S. Mossa, A. Turtschi, and L. A. N. Amaral, The worldwide air transportation network: Anomalous centrality, community structure, and cities’ global roles, Proceedings of the National Academy of Sciences 102, 7794 (2005).

[58] R. Miralles, P. J. Gerrish, A. Moya, and S. F. Elena, Clonal interference and the evolution of rna viruses, Science 285, 1745 (1999).

[59] J. Arjan G, M. d. Visser, C. W. Zeyl, P. J. Gerrish, J. L. Blanchard, and R. E. Lenski, Diminishing returns from mutation supply rate in asexual populations, Science 283, 404 (1999).

[60] N. Colegrave, Sex releases the speed limit on evolution, Nature 420, 664 (2002).

[61] A. C. Shaver, P. G. Dombrowski, J. Y. Sweeney, T. Treis, R. M. Zappala, and P. D. Sniegowski, Fitness evolution and the rise of mutator alleles in experimental escherichia coli populations, Genetics 162, 557 (2002).

[62] D. E. Rozen, J. A. G. De Visser, and P. J. Gerrish, Fitness effects of fixed beneficial mutations in microbial populations, Current biology 12, 1040 (2002).

[63] K. C. Kao and G. Sherlock, Molecular characterization of clonal interference during adaptive evolution in asexual populations of saccharomyces cerevisiae, Nature Genetics 40, 1499 (2008).

[64] A. Hagberg, P. J. Swart, and D. A. Schult, Exploring network structure, dynamics, and function using NetworkX, Tech. Rep. (Los Alamos National Laboratory (LANL), Los Alamos, NM (United States), 2008).

